# Purine biosynthesis pathways are required for myogenesis in *Xenopus laevis*

**DOI:** 10.1101/2023.08.17.553788

**Authors:** Maëlle Duperray, Elodie Henriet, Christelle Saint-Marc, Eric Boué-Grabot, Bertrand Daignan-Fornier, Karine Massé, Benoît Pinson

## Abstract

Purines are required for fundamental biological processes and alterations in their metabolism lead to severe genetic diseases associated with developmental defects whose etiology remains unclear. Here, we studied the developmental requirements for purine metabolism using the amphibian *Xenopus laevis* as a vertebrate model. We provide the first functional characterization of purine pathway-genes and show that these genes are mostly expressed in nervous and muscular embryonic tissues. Morphants were generated to decipher the functions of these genes, with a focus on the adenylosuccinate lyase (*ADSL*), an enzyme required for both salvage and *de novo* purine pathways. *adsl.L* knock-down leads to severe reduction in the expression of the Myogenic Regulatory Factors (MRFs: Myod1, Myf5 and Myogenin), thus resulting in defects in somitogenesis and in the formation and/or migration of both craniofacial and hypaxial muscle progenitors. Similar alterations were observed upon reduced expression of *hprt1.L* and *ppat*, two genes specific to the salvage and the *de novo* pathways, respectively. In conclusion, our data shows for the first time that *de novo* and recycling purine pathways are essential for myogenesis and highlight new mechanisms in the regulation of MRFs gene expression.

## 1. Introduction

Purine triphosphate nucleotides, ATP and GTP are synthesized via a highly conserved [1] *de novo* pathway allowing sequential construction of the purine ring on a ribose phosphate moiety provided by phosphoribosyl pyrophosphate (PRPP) (Figure 1A and Figure S1 for details). This *de novo* pathway results in the synthesis of inosine 5’-monophosphate (IMP) that can be converted into either AMP or GMP and then into other phosphorylated nucleotide forms required for all known forms of life. These nucleotides can alternatively be synthetized *via* a salvage pathway using precursors taken up from the extracellular medium or coming from the internal recycling of preexisting purines (Figure 1A). Purines and their derivatives (such as NAD(P) (nicotinamide adenine dinucleotide (phosphate)), FAD (Flavin adenine dinucleotide), coenzyme A, ADP-ribose, S-adenosyl-methionine and S-adenosyl-homocysteine) form a family of metabolites among the most abundant (mM range) in cells and are involved in a myriad of physiological events. Purines are required for numerous cellular processes such as biosynthesis of nucleic acids and lipids, replication, transcription, translation, maintenance of energy and redox balances, regulation of gene expression (methylation, acetylation…) and cell signaling, such as the purinergic signaling pathway [2]. Therefore, defects, even minor, in the purine *de novo* biosynthesis or recycling pathways lead to deleterious physiological effects. Indeed, to date, no less than 35 rare genetic pathologies have been associated with purine metabolism dysfunctions [3–7].

**Figure 1.**
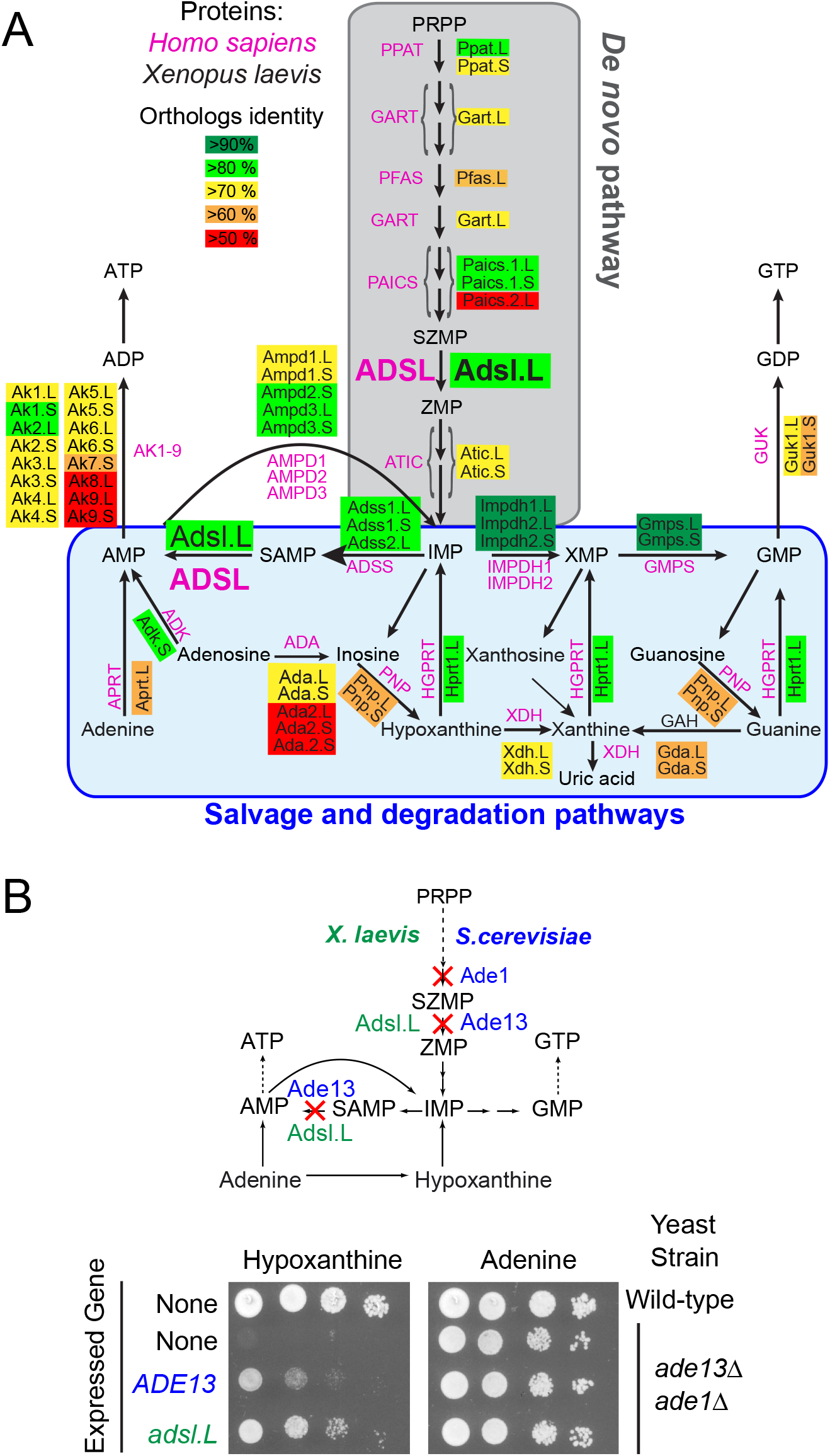
The *X. laevis adsl.L* gene encodes the adenylosuccinate lyase activity required in two non-sequential steps of the highly conserved purine synthesis pathways. (**A**) Schematic representation of the human and *X. laevis* purine biosynthesis pathways. Abbreviations: AMP: adenosine monophosphate; GMP: guanosine monophosphate; IMP: Inosine monophosphate; PRPP: Phosphorybosyl pyrophosphate; SAMP: Succinyl-AMP; SZMP: Succinyl-Amino Imidazole Carboxamide Ribonucleotide monophosphate; XMP: Xanthosine monophosphate; ZMP: Amino Imidazole CarboxAmide Ribonucleotide monophosphate. (**B**) Functional complementation of the growth defect of the yeast adenylosuccinate lyase deletion mutant by expression of the *X. laevis adsl.L* ortholog gene. Yeast wild-type and *adsl* knock-out mutant (*ade13 ade1*) strains were either transformed with a plasmid allowing expression of the *Saccharomyces cerevisiae* (*ADE13*) or the *X. laevis* (*adsl.L*) adenylosuccinate lyase encoding genes, or with the empty vector (None). Serial dilutions (1/10) of transformants were dropped on SDcasaWA medium supplemented with either hypoxanthine or adenine as the sole external purine source. Plates were incubated for 48 h at 37 °C before imaging. Of note the *ade1 ade13* double deletion mutant was used in this experiment to avoid genetic instability associated with the accumulation of SZMP and/or its nucleoside derivatives observed in the single *ade13* single mutant [22].

Purine-associated pathologies share a broad spectrum of clinical symptoms including hyperuricemia, severe muscular and neurological dysfunctions and often immunological, hematological and renal manifestations [3,4]. They are all characterized by abnormal levels of purine nucleotides, nucleosides and/or nucleobases in the patient’s body fluids and/or cells. Although in most cases the mutated gene has been identified, the causal link between the defective enzyme, the decrease in some final products, the accumulation of purine intermediates, or the alteration of physiological functions with the observed symptoms often remains very elusive, even unknown.

The adenylosuccinate lyase deficiency (OMIM 103050) is one of the most studied purine metabolism pathologies and alters the Adsl enzyme, that catalyzes two non-consecutive reactions required both for the salvage and *de novo* purine pathways (Figure 1A). First described in 1984 [8], this rare autosomal recessive disorder is associated with a massive accumulation in patient fluids (blood, urine and cerebrospinal fluid) of succinyl derivatives (Succinyl-adenosine and Succinyl-AICAR (AminoImidazole CarboxAmide Ribonucleoside)) corresponding to dephosphorylation of the monophosphate substrates of Adsl (SZMP and SAMP, Figure 1A). The first patient-specific mutation in *ADSL* gene was identified in 1992 [9] and, to date, more than 50 different mutations have been described [10] (Figure S2, orange and yellow boxes). All the biochemically-studied mutations lead to a low residual Adsl enzymatic activity [11–15]. This pathology is characterized by serious neuro-muscular dysfunctions including psychomotor retardation, brain abnormalities, autistic features, seizures, ataxia, axial hypotonia, peripheral hypotonicity, muscular wasting and growth retardation [13,16,17]. Three distinct forms of *ADSL* deficiency have been categorized based on the severity of the symptoms: a fatal neonatal form (respiratory failure causing the death within the first weeks of life), a childhood form with severe neuromuscular symptoms (severe form; type I) and a more progressive form with milder symptoms (mild form; type II) (for review see [18]). A robust correlation has been clearly established between the residual activity of Adsl, the consequent accumulation of succinyl derivatives and the intensity of phenotypes [15,19–21]. However, these studies do not explain the molecular bases associated with *ADSL* deficiency, though they raise a real interest in identifying biological functions that could be modulated by SZMP, SAMP and their derivatives. For example, SZMP acts as a signal metabolite regulating transcriptional expression in yeast [22], it promotes proliferation in cancer cells by modulating pyruvate kinase activity [23] and SAMP stimulates insulin secretion in pancreatic cells [24]. More recently, RNA-seq analysis of a cellular model of ADSL deficiency identified misexpression of genes involved in cancer and embryogenesis, bringing some first clues to understanding the molecular bases of this rare disease [25].

To go further in the etiology of this disease, it is now essential to identify the different biological processes that are altered by a purine deficiency during embryonic development as the most severe symptoms appear *in utero* or within the first weeks or months after birth. To our knowledge, few invertebrate and vertebrate models have been developed to study this disease and no mammalian model recapitulating *ADSL* deficiency exists, most likely due to embryonic lethality [26,27]. Here, we set up a *X. laevis* model to assess the developmental functions of *ADSL* by knocking down its embryonic expression. This powerful vertebrate model organism has been widely used to identify developmental impairments associated with human pathologies [28,29].

We report the identification and functional validation of the *adsl.L* gene and show this gene as being mostly expressed in the nervous and muscular tissues during *X. laevis* development. Knock-down of *adsl.L* results in down-regulation at different developmental stages of several myogenic regulating factors (MRF), such as Myod1, Myf5 and Myogenin, expression and leads to alterations in somites, craniofacial and hypaxial muscle formation, thus showing that *adsl.L* is essential for myogenesis. Finally, functional comparative analysis of other enzymes in the *de novo* and salvage purine pathways (Ppat.L, Ppat.S, and Hprt1.L) highlights a major role for purine metabolism in muscle tissue development, providing insight into the developmental defects that underlie some of the symptoms of patients with severe ADSL deficiency as well as other purine-associated pathologies.

## 2. Materials and Methods

### 2.1 Yeast media

SDcasaW is SD medium (0.5 % ammonium sulfate, 0.17 % yeast nitrogen base without amino acids and ammonium sulfate (BD-Difco; Franklin Lakes, NJ, USA) and 2 % glucose) supplemented with 0.2% casamino acids ((#A1404HA; Biokar/Solabia group; Pantin, France) and tryptophan (0.2 mM). When indicated, adenine (0.3 mM) or hypoxanthine (0.3 mM) was added as an external purine source in SDcasaW.

### 2.2 Yeast strains and plasmids

All yeast strains are listed in Table S1 and belong to, or are derived from, a set of disrupted strains isogenic to BY4741 or BY4742 purchased from Euroscarf (Germany). Double mutant strains were obtained by crossing, sporulation and micromanipulation of meiosis progeny. All plasmids (Table S2) were constructed using the pCM189 vector [30], allowing expression of *X. laevis* gene in yeast under the control of a tetracycline-repressible promoter. *X. laevis* open reading frame (ORF) sequences were amplified by PCR from I.M.A.G.E Clones (Source Biosciences; Nottingham, UK). All *X. laevis* ORF sequences were fully verified by sequencing after cloning in the pCM189 vector. Further cloning details are available upon request.

### 2.3 Yeast growth test

For drop tests, yeast transformants were pre-cultured overnight on solid SDcasaWA medium, re-suspended in sterile water at 1−10^7^ cells/ml and submitted to 1/10 serial dilutions. Drops (5 μl) of each dilution were spotted on freshly prepared SDcasaW medium plates supplemented or not with adenine or hypoxanthine. Plates were incubated either at 30 or 37 °C for 2 to 7 days before imaging.

### 2.4 Embryo culture

*X. laevis* males and females were purchased from the CNRS Xenopus Breeding Center (CRB, Rennes, France). Embryos were obtained by *in vitro* fertilization of oocytes collected in 1X Marc’s Modified Ringers saline solution (1X MMR:100 mM NaCl, 2 mM KCl, 2 mM CaCl_2_, 1 mM MgSO_4_, 5 mM Hepes, pH 7.4), from a hormonally (hCG (Agripharm), 750 units) stimulated female by adding crushed testis isolated from a sacrificed male. Fertilized eggs were de-jellied in 3% L-cysteine hydrochloride, pH 7.6 (Sigma-Aldrich), and washed several times with 0.1X MMR. Embryos were then cultured to the required stage in 0.1X MMR in the presence of 50 µM of gentamycin sulfate. Embryos were staged according to the Nieuwkoop and Faber table of *X. laevis* development [31].

### 2.5 mRNA synthesis and morpholino oligonucleotides

Capped mRNAs were synthesized using Sp6 mMESSAGE mMACHINE Kits (Ambion) from linearized plasmids (listed in Table S3). *Adsl.L*, *ppat.S*, *ppat.L* and *hprt1.L* RNA were transcribed from the IMAGE clones. *Adsl.L*-RNA*, *ppat.S*-RNA*, *ppat.L*-RNA* and *hprt1.L*\* ORFs were amplified by PCR in conditions which allow the introduction of mutations in the morpholino oligonucleotide (MO) binding site and were subcloned into the plasmid pBF. *Homo sapiens ADSL cDNA was subcloned into the plasmid pCS2+. Adsl.L* MO1 (5’-AAGCATGGAGGGGAGCAGTGGGCTAAG-3’), *Adsl.L* MO2 (5’-ATGGAGGGGAGCAGTGGGCTAAGCAT-3’), *hprt1.L* MO (5’-GGACACAGGCTCAGACATGGCGAGC-3’), *ppat.L*/*ppat.S* MO (5’-GTGATGGAGTTTGAGGAGCTGGGGAT-3’) and standard control MO (cMO) were designed and supplied by Gene Tools, LLC. The position of the MOs in relation to their respective RNA is indicated in Figure S3.

### 2.6 Microinjections

Embryos were injected with MO alone or in combination with MO non-targeted mRNA (mutated *Adsl.L* RNA* or *Homo sapiens ADSL*, as specified in the text/legend) into the marginal zone of one blastomere at the 2-cell stage. *LacZ* (250 pg) RNA was co-injected as a lineage tracer. Embryos were injected in 5% Ficoll, 0.375X MMR, cultured to various developmental stages, fixed in MEMFA (MOPS 100 mM pH 7.4, EGTA 2 mM, MgSO_4_ 1 mM, 4.0 % (v/v) formaldehyde) and stained for beta-galactosidase activity using Red-Gal or X-Gal substrates (RES1364C-A103X or B4252, Merck) to identify the injected side and correctly targeted embryos. Embryos were fixed again in MEMFA before dehydration into 100% methanol or ethanol for immunohistochemistry or *in situ* hybridization, respectively.

### 2.7 In situ hybridization

Whole-mount *in situ* hybridizations (ISH) were carried out as previously described [32–34]. Sense and antisense riboprobes were designed by subcloning fragments of coding cDNA sequences in pBlueScript II (SK or KS) plasmids (Addgene). Riboprobes were generated by *in vitro* transcription using the SP6/T7 DIG RNA labelling kit (Roche, # 11175025910) after plasmid linearization as indicated in Table S4. Riboprobe hybridization detection was carried out with an anti-DIG Alkaline Phosphatase antibody (Roche, #11093274910) and the BM-Purple AP substrate (Roche, #11442074001). Riboprobes for *lbx1*, *myf5*, *myod1*, *myogenin*, *pax3* and *tbxt* (*xbra*) were previously described [35–37].

### 2.8 Immunostaining

Whole-mount immunostaining of differentiated skeletal muscle cells was performed using the monoclonal hybridoma 12/101 primary antibody [38] (1/200 dilution; DSHB #AB-531892) and the EnVision^+^ Mouse HRP kit (Agilent Technologies, K4007) according to the manufacturer’s recommendations.

### 2.9 Temporal expression of genes established by RT-PCR

RNA extraction from whole embryos, cDNA synthesis and RT-PCR were performed as previously described [33,39]. Sequences of the specific primers designed for each gene and PCR amplification conditions are given in Table S5. Chosen primers were selected to differentiate homeolog gene expression and to discriminate potential genomic amplification from cDNA amplification. PCR products were verified by sequencing (Eurofins genomics). The quantity of input cDNA was determined by normalization of the samples with the constant ornithine decarboxylase gene *odc1.L* [40]. Linearity of the signal was controlled by carrying out PCR reactions on doubling dilutions of cDNA and negative controls without either RNAs, reverse transcriptase and cDNA were also performed. Experiments were done at least twice (N ≥ 2) on embryos from two different females and representative profiles are shown in Figure 2.

**Figure 2.**
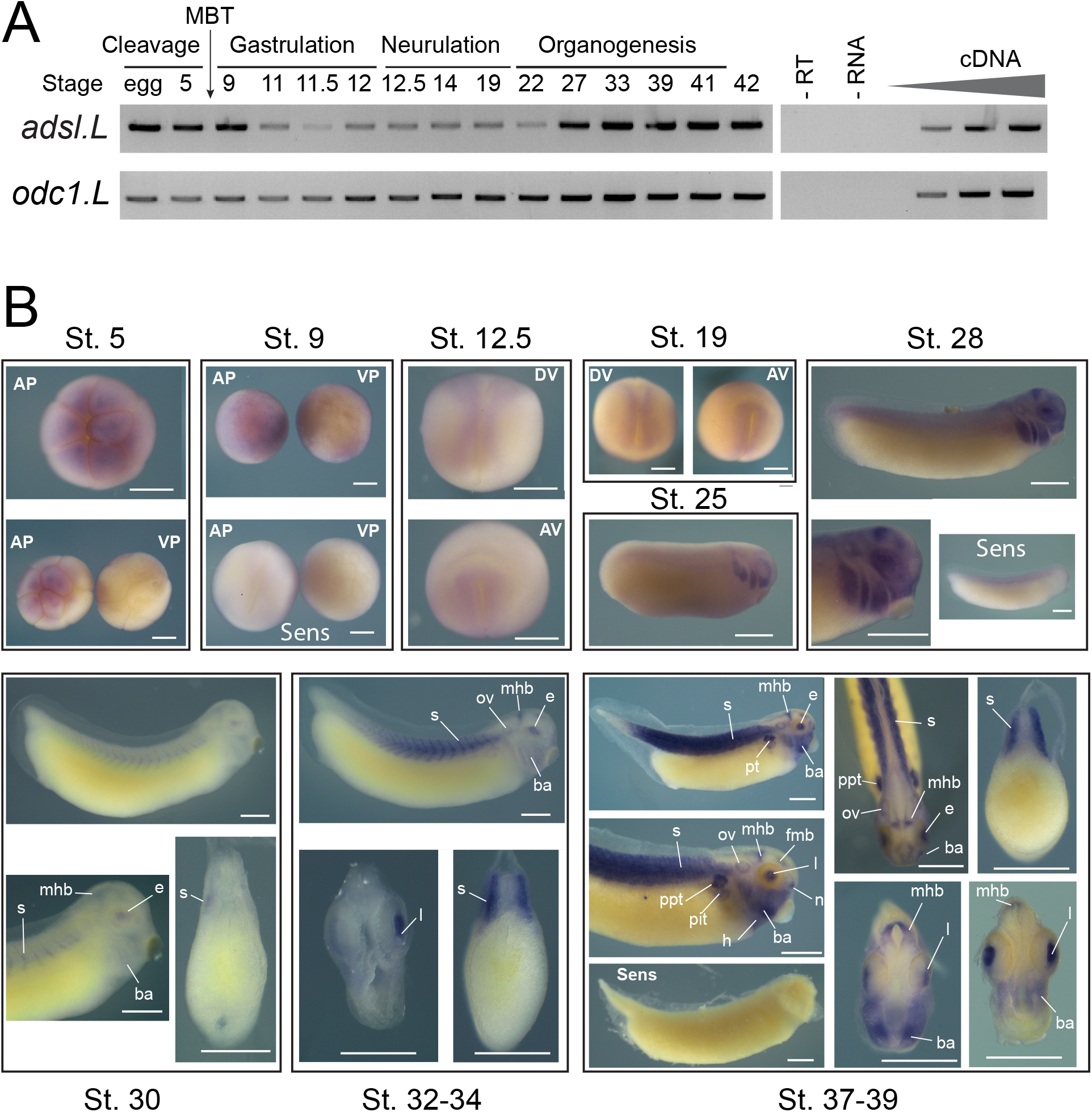
Spatiotemporal expression of *adsl.L* gene during *X. laevis* embryonic development. (**A**) Temporal expression profiles of *adsl.L* gene during embryogenesis. The expression profile was determined by RT-PCR from the cDNA of the fertilized oocyte (egg) and whole embryo at indicated stages covering the different phases of *X. laevis* embryogenesis. The ornithine decarboxylase gene *odc1.L* was used as a loading control and negative controls were performed by omitting either reverse transcriptase (-RT) or RNA (-RNA) in the reaction mix. Linearity was determined by dilutions of cDNA from stage 39 for *adsl.L* and 41 for *odc1.L*. Mid-blastula transition (MBT) is indicated. (**B**) Spatial expression profile of *adsl.L* gene during embryogenesis. Whole-mount *in situ* hybridization with *adsl.L*-specific DIG-labelled antisense or sense RNA probes was performed on embryos from stages (St.) 5 to 37/39. St. 5 and 9: animal (AP) and vegetal (VP) pole views, St. 12.5 and 19: dorsal (DV) and anterior (AV) views; later stages: lateral views, with dorsal up and anterior on the right, and dorsal view at stage 37-39. Transverse sections are dorsal up. Abbreviations: ba, branchial arches; e, eye; fmb, forebrain**-**midbrain boundary; h, heart; l, lens; mhb, midbrain-hindbrain boundary; n: nasal placode; ov, otic vesicle; ppt, pronephric proximal tubules; pit, pronephric intermediate tubules; s, somites. Bars: 0.5 mm.

### 2.10 Embryo scoring and photography

Embryos were bleached (1% H_2_O_2,_ 5% formamide, 0.5X SSC) to remove all visible pigment and phenotypes were determined in a commonly used way, blind-coded, by comparing the injected and un-injected sides. Only embryos with normal muscle tissue formation on the un-injected side and correctly targeted β-galactosidase staining on the injected side were scored. Transverse sections were performed with a razor blade on fixed embryos. Embryos were photographed using a SMZ18 binocular system (Nikon).

### 2.11 Statistics and reproducibility

All experiments were carried out on at least two batches of fertilized eggs from two independent females. Histograms represent the percentage of embryos displaying each phenotype and the number of embryos in each category is indicated. Fisher’s exact test was used for statistical analyses and p-values are presented above the bars of the histogram in all figures.

### 2.12 Bioinformatics

Sequences were identified on the NCBI and Xenbase databases [41]. Basic Local Alignment Search Tool (BLAST) searches were performed on the NBCI Nucleotide and the Xenbase *X. laevis* 9.1 Scaffolds genome databases [42]. Conceptual translation of complementary DNA (cDNA) was performed on the ExPASy Internet website using the program Translate Tool (web.expasy.org/translate/). Comparison of coding sequences from *H. sapiens*, *S. cerevisiae*, *X. laevis* and *X. tropicalis* were done on https://blast.ncbi.nlm.nih.gov. Accession numbers of all sequences used in this study are given in Tables S6 and S7.

### 2.13 Ethics statement

All experimental procedures, performed in the Aquatic facility of the Centre Broca-Nouvelle Aquitaine in Bordeaux EU0655, complied with the official European guidelines for the care and use of laboratory animals (Directive 2010/63/EU) and were approved by the Bordeaux Ethics Committee (CEEA50) and the French Ministry of Agriculture (agreement #A33 063 942, approved 2018/07/27).

## 3. Results

### 3.1 Identification of X. laevis purine pathway genes

The purine biosynthesis pathways are known to be highly conserved throughout evolution [1]. Most of *X. laevis* genes encoding potential purine-pathway enzymes have been putatively identified and annotated by automated computational analyses from the entire genome sequencing [43]. Protein sequences were deduced from conceptual translation of these annotated genes and aligned with their potential orthologs, e.g. *H. sapiens* and *X. tropicalis* (Table S6). The vast majority of *X. laevis de novo* and salvage pathways have two homeologs, whose protein sequences display a high degree of identity with their *X. tropicalis* and human orthologs (more than 85 and 50 %, respectively) (Figure 1A and Table S6), suggesting that these annotated genes effectively encode the predicted purine-pathway enzymes.

To establish that the putative *X. laevis* purine-pathway genes encode the predicted enzymatic activities, a functional complementation assay was undertaken by heterologous expression of *X. laevis* genes in the yeast *Saccharomyces cerevisiae,* which has been proven to be a valuable model to investigate metabolic pathways [44]. Yeast knock-out mutants were available in the laboratory and sequence alignments showed a high degree of identity between *S. cerevisiae* and *X. laevis* orthologous protein sequences (Figure S1 and Table S7). Plasmids-allowing expression of the *X. laevis* ORF in *S. cerevisiae* were transformed in the cognate yeast knock-out mutant, and functional complementation was tested. As shown in Figure 1B, the yeast *adsl* knock-out mutant (*ade13*) was unable to grow in the presence of hypoxanthine as a unique purine source, but growth was restored by the expression of either the *S. cerevisiae ADE13* or the *X. laevis adsl.L* gene. By contrast, all these strains were able to grow in the presence of adenine that allow purine synthesis *via* the adenine phosphoribosyl transferase (Apt1) and AMP deaminase (Amd1) activities (Figure S1). Similar experiments were conducted with 15 other *X. laevis* purine-pathway encoding genes, 13 of which were able to complement the *S. cerevisiae* cognate knock-out mutants (Figure S4 and Figure S5). Altogether, this data allowed us to functionally validate *adsl.L* and 13 other *X. laevis* genes involved in purine *de novo* and recycling pathways, demonstrating that the purine pathways are functionally conserved in *X. laevis*.

We then focused on the role of adenylosuccinate lyase as: (1) it is encoded by a single gene (*adsl.L)*, greatly simplifying knock-down experiments, and its protein sequence is highly conserved between human and *X. laevis* (Figure S2), (2) it is the only enzymatic activity that is required in both the *de novo* biosynthesis and recycling purine pathways, and (3) its mutation leads to severe developmental alterations, for which the molecular bases, associated with the symptoms, are still largely unknown.

### 3.2 The adsl.L gene is mostly expressed in muscular and neuronal tissues and their precursors during X. laevis development

*Adsl.L* spatiotemporal expression during *X. laevis* development was established by two complementary approaches. First, its temporal expression profile was determined by RT-PCR on whole embryos from fertilized egg to stage 45 (Figure 2A). The *adsl.L* gene displays both a maternal (before the mid-blastula transition (MBT)) and a zygotic expression at all developmental stages studied.

*Adsl.L* spatial expression was then determined by *in situ* hybridization (Figure 2B). *Adsl.L* expression is detected in the animal, but not vegetal, pole during the cleavage phase. Zygotic expression was detected in the neuroectoderm and in the paraxial mesoderm during neurulation. From stage 25 until the late organogenesis stages, *Adsl.L* expression was found in the developing epibranchial placodes, lens and in the central nervous system (fore-midbrain and mid-hindbrain boundaries). From stage 30 onwards, *Adsl.L* transcripts were also detected in the somites and its somitic expression appeared to increase during organogenesis. At the late organogenesis stage, its expression was detected in other mesoderm-derived tissues, such as the pronephric tubules (proximal and intermediate tubules) and the heart.

In parallel, using similar approaches, we showed that (1) 14 out of the 16 tested genes display a similar temporal expression profile to that of *adsl.L* (Figure S6) and (2) all 9 genes tested by *in situ* hybridization are expressed in neuro-muscular tissues (e.g. somites, hypaxial muscles, central nervous system and eyes), even if differences in the tissue specificity exist for some of these genes (Figures S7-S8). This is consistent with the fact that neuromuscular dysfunctions are the primary symptoms associated with purine-dependent diseases.

### 3.3 The adsl.L gene is required for X. laevis myogenesis

To understand the role of *Adsl.L* during muscle development, we have undertaken a loss of function approach using two specific anti-sense morpholino oligonucleotides (MOs) (see Figure S3A for MOs efficiency). MOs injections were performed in the marginal zone of one blastomere at the 2-cell stage. This injection has the advantage of unilaterally affecting the development of the future mesoderm, with the non-injected side serving as an internal control. A dose-dependent curvature along the anterior/posterior axis was observed following injection of *adsl.L* MO1 and MO2, and not of the control morpholino (cMO), with a concave deformation always corresponding to the side of injection (Figure 3A-B). This curvature phenotype was already documented when myogenesis was altered [45], suggesting that *adsl.L* gene could be required for proper myogenesis in *X. laevis*. To test this hypothesis, we performed immunostaining on tailbud and tadpole stages with the differentiated skeletal muscle cells-specific 12-101 antibody [38]. cMO injected embryos displayed normal global morphology and normal muscle phenotype at all stages tested. By contrast, myotomes were smaller along the dorsoventral axis following the injection of 10 ng of *adsl.L* MO1 or MO2 at the tailbud stage (Figure 3 C-D). Furthermore, somite morphology were significantly altered. Straight-shape somites, blurred somite domains and fewer v-shape somites were observed on the injected side. 12-101 positive area was also reduced along the anteroposterior axis, the latter shortening most likely being at the origin of the curvature phenotype observed (Figure 3A). This muscle alteration was even more exacerbated by increasing the dose of MO, with severe reduction, and in some cases, absence of 12/101 positive domain and no somitic-like structure observed on the injected side in more than 80% of the injected 20 ng MO1 embryos (Figure 3C). This dose-dependent muscle phenotype was still observed in the tadpole stage (stage 39) with a similar percentage of *Adsl.L* morphant embryos displaying myotome reduction and altered chevron-shaped somites in their injected side, especially in the anterior trunk region (Figure S9A-B). At the late tadpole stage (stage 41), somite defects, characterized by anterior straight or criss-crossed somites, were observed on the injected side of *adsl.L* morphants (red arrowheads; Figure 3E-F), whereas the cMO embryo injected side displayed v-shaped somites (white arrowheads; Figure 3E-F). Reduction of the hypaxial muscles (precursors of the limb muscles) was also observed in *adsl.L* morphants but not in cMO injected embryos (Figure 3E, G, compare the blue to the white dashed lane in *adsl.L* MO1 embryo). This hypaxial muscle phenotype was worsened when the MO dose was increased with no hypaxial muscle progenitors in 70% and 96% of analyzed embryos following 10 ng and 20 ng of *Adsl.L* MO1 injection respectively (Figure S9A, C). These phenotypes were rescued by co-injecting the *adsl.L* mutated RNA*, whose translation is unaffected by *adsl.L* MO1 and MO2 (Figure S3A), with *adsl.L* MO1 (Figure 3E-G). No significant phenotype was obtained by co-injecting the *adsl.L* RNA* with cMO (Figure 3E-G). Together, these results validate the muscular-associated phenotypes as a consequence of a specific *adsl.L* gene knock-down and not of the potential morpholino injection’s side effects.

**Figure 3.**
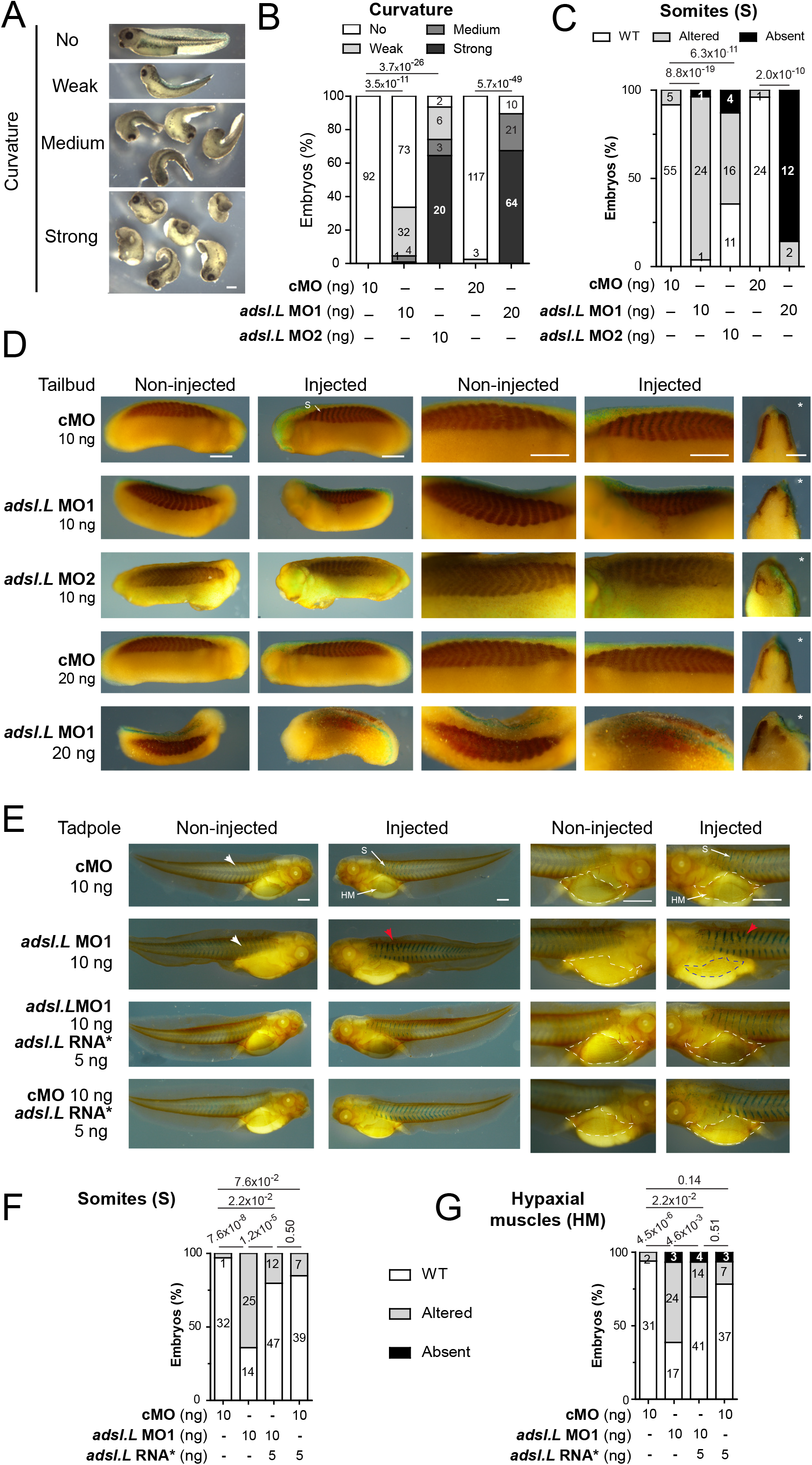
The *adsl.L* gene is required for somites and hypaxial muscle formation in *X. laevis*. (**A**) Representative images of *adsl.L* morphants following *adsl.L* MO1 injection at either 10 ng (weak and medium curvature) or 20 ng (medium and strong curvature). (**B**) Quantification and statistics of the curvature phenotype. (**C-G**) Immunostaining with the differentiated muscle cells specific 12-101 antibody revealed a strong alteration of somites and hypaxial muscle formation in *adsl.L* knock-down and *adsl.L* over-expressing embryos. Representative images (**D-E**), quantification and statistics (**C**, **F-G**) of somites and hypaxial muscles phenotype at tailbud (**C**-**D**) and tadpole (**E**-**G**) stages. Injected side is indicated by asterisks. S: somites; HM: Hypaxial muscles. Bars: 0.5 mm. White and red arrowheads point to typical somite chevron shapes and altered somites, respectively. The a*dsl.L* RNA* refers to mutated RNA whose translation is not affected by the *adsl.L* specific MOs (Figure S3). Numbers above the bars of histograms correspond to p-values.

To determine whether the observed muscle phenotypes were specifically related to the knock-down of the *adsl.L* gene or to the purine pathways alteration, similar loss of function experiments (see Figure S3B-C for MOs efficiency) were carried out on other purine-associated genes: (1) the *ppat* (*ppat.L* and *ppat.S*) genes encoding the phosphoribosylpyrophosphate amidotransferase (Ppat), the first enzyme of the purine *de novo* pathway, and (2) the *hprt1.L* gene encoding the Hypoxanthine-PhosphoRybosylTransferase (Hprt) in the salvage pathway (Figure 1A), the enzyme allowing metabolization of hypoxanthine, the major purine source in *X. laevis* embryos [46]. Comparable muscle alterations were obtained with independent knock-down of these genes, *i.e.* a strong alteration of somite shape at both the tailbud and tadpole stages, associated with smaller myotomes at the early organogenesis stage and reduction, or even absence, of hypaxial muscles in the latter stages (Figure S10 and S11). Taken together, these similar muscle phenotypes observed in *adsl.L*, *ppat* and *hrpt1.L* morphants demonstrate a strong association between altered purine pathways and defects in somitogenesis and myogenesis.

### 3.4 Expression of the myod1 and myf5 genes requires functional purine pathways

We assessed whether the muscular phenotypes observed upon knock-down of purine-associated genes could result from an alteration of the myogenic regulatory factor’s (MRFs) expression during the different myogenic waves [36,47]. Expression of both *myod1* and *myf5* was first analyzed by *in situ* hybridization at the early neurula stage, during the first myogenic wave (Figure 4 A-D). A strong reduction of both *myod1* and *myf5* expression was observed on the injected side of *adsl.L* morphants (Figure 4 A, C), whereas no alteration could be seen in cMO-injected embryos. This significant reduction was even stronger in the anterior region, which will give rise to the first somites. Importantly, *myod1* expression was rescued by the injection of *X. laevis adsl.L* mutated RNA* or human *ADSL* RNA in *adsl.L* MO1 and MO2 morphants (Figure 4A-B). These results are consistent with an adenylosuccinate lyase activity, even heterologous, being required for the proper expression of *myod1*. Of note, over-expression of either *X. laevis adsl.L* mutated RNA* or human *ADSL* mRNA induced a small but significant increase in *myod1* expression (Figure S12). To exclude that the observed reduction of *myod1* and *myf5* expression resulted from an alteration of mesoderm formation and/or integrity in *adsl.L* morphants, expression of the pan-mesoderm marker *tbxt* (*xbra*) was analyzed by *in situ* hybridization on whole embryos at stage 11. No significant difference was observed in all injected embryos (Figure 4E). Finally, a strong and similar reduction in *myod1* expression, with no alteration of the *tbxt* gene expression pattern, was also observed in either *ppat.L/ppat.S* or *hprt1.L* morphants (Figure S13). Altogether, these results show that functional purine synthesis is strictly required for a proper expression of the myogenic regulatory factor genes *myod1* and *myf5* during the first myogenic wave.

**Figure 4.**
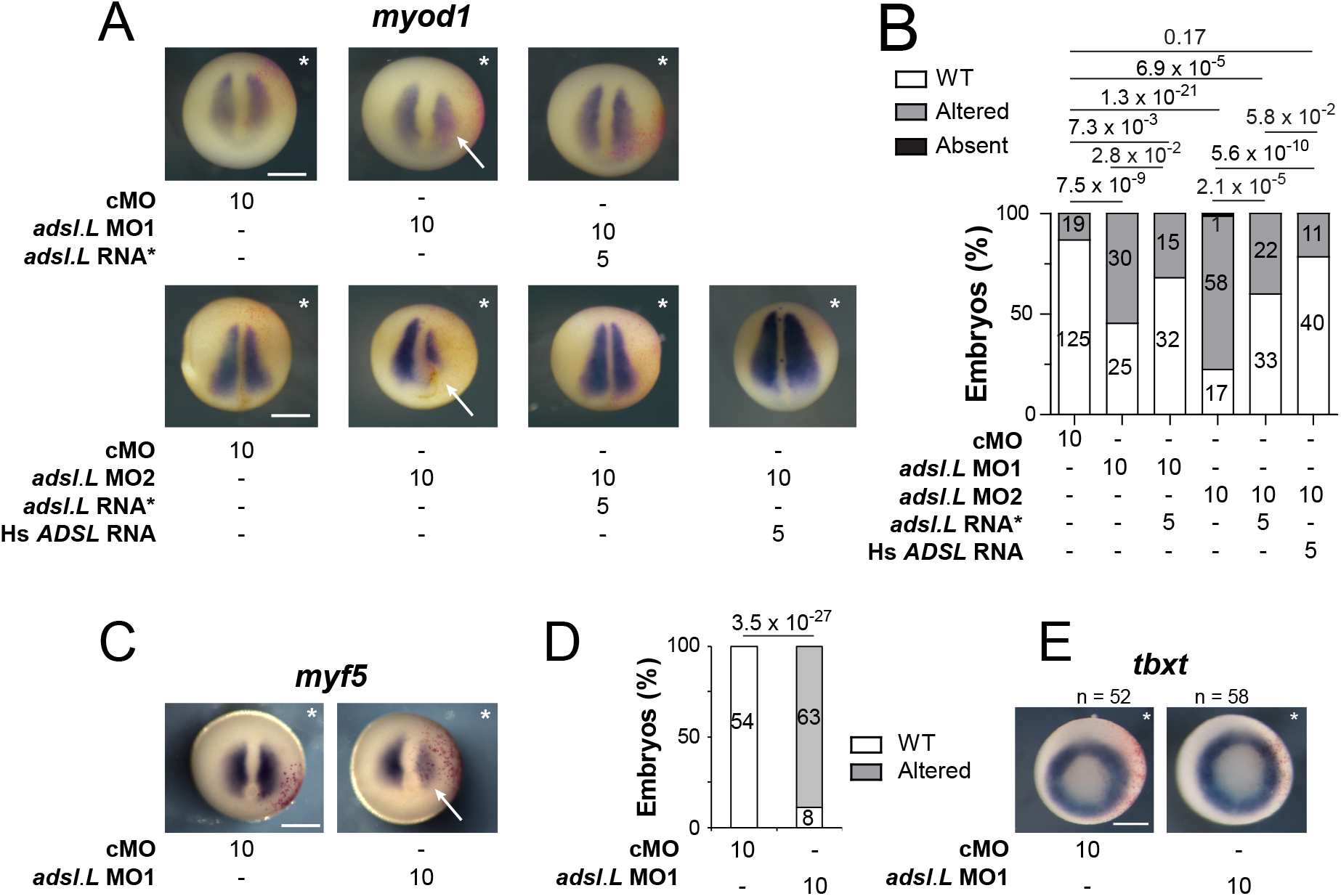
Expression of the myogenic regulatory factors *myod1* and *myf5* in paraxial mesoderm is strongly affected by the knock-down of the *adsl.L* gene. (**A**) Expression of *myod1* gene at the early neurula stage is altered by the knock-down of *adsl.L* gene and rescued by *adsl.L* RNA* and human *ADSL* RNA. (**C**) Representative images of the effect on *myf5* RNA expression by *adsl.L* knock-down at stage 12.5 revealed by *in situ* hybridization. (**B, D**) Quantification and statistics of the *myod1* and *myf5* expression phenotypes are presented in (**A**) and (**C**), respectively. (**E**) Knock-down of *adsl.L* does not alter mesoderm formation, as revealed by the absence of change in *tbxt* (*xbra*) RNA expression domain at stage 11. Injected side is indicated by asterisks. The amounts of MO and RNA presented in this figure are in ng. Bars: 0.5 mm. Numbers above the bars of histograms correspond to p-values.

To further analyze the effects of *adsl.L* knock-down on MRF expression, *myod1* and *myf5* expression was then analyzed at later developmental stages (Figure 5). At the tailbud stage, the *myod1* expression in the somites was found significantly altered in *adsl.L* MO1 and MO2 morphants. As observed with the 12/101 staining, myotomes were shortened along the dorsoventral axis and somite shape defects, and even loss of the chevron shape of the most anterior somites, were observed (blue arrowheads, Fig 5 A-B). *Myod1* expression in the ventral border of the dermomyotome was also significantly reduced or lost (red arrowheads, Fig 5 A-B), while its expression in the dermomyotome dorsal border was less altered in the anterior trunk region. Expression of *myod1* in craniofacial muscles was also reduced, or even absent following *adsl.L* MO1 or MO2 injection (green arrowheads, Fig 5 A-B). These alterations in craniofacial muscles were also observed at the tadpole stages with a severe significant reduction of *myod1* expression especially in the interhyoideus (ih), quadrato-hyoangularis and orbitohyoideus anlagen (q/oh), in *adsl.L* MO1 and MO2 morphants (light green arrowheads, Fig 5C-D). Expression of *myf5* was also significantly altered in craniofacial muscles at both the tailbud and tadpole stages, even if these alterations are milder than the *myod1* ones (Figure 5E-H). Indeed, *myf5* expression in *adsl.L* morphants is significantly reduced or absent at both stages in the dorsal region of the pharyngeal arch muscle anlagen (black arrowhead, Figure 5G), whereas its expression in the interhyoideus and the intermandibularis muscle anlagen is rarely reduced (pink arrowhead, Figure 5G). However, no alteration of *myf5* expression was observed in the undifferentiated pre-somitic mesoderm in the tail region (yellow circles, Figure 5E-H). The implication of *adsl.L* in the formation of the cranial muscles was also confirmed by analyzing the expression of myogenin (*myog*) following *adsl.L* MO1 and MO2 injection at stage 39 (Figure 6A and Figure S14). A strong reduction, even an absence, of *myogenin* staining in the pharyngeal arch muscle anlagen was observed on the injected side in *adsl.L* morphants, whereas its expression in the interhyoideus and intermandibularis muscle anlagen was, however, less affected. These results are consistent with those observed by 12-101 immunolabelling on *adsl.L* morphants (Figure S9, blue arrowhead). Altogether, these data show that *adsl.L* is required for *myod1*, *myf5* and *myogenin* expression during the second wave of myogenesis.

**Figure 5.**
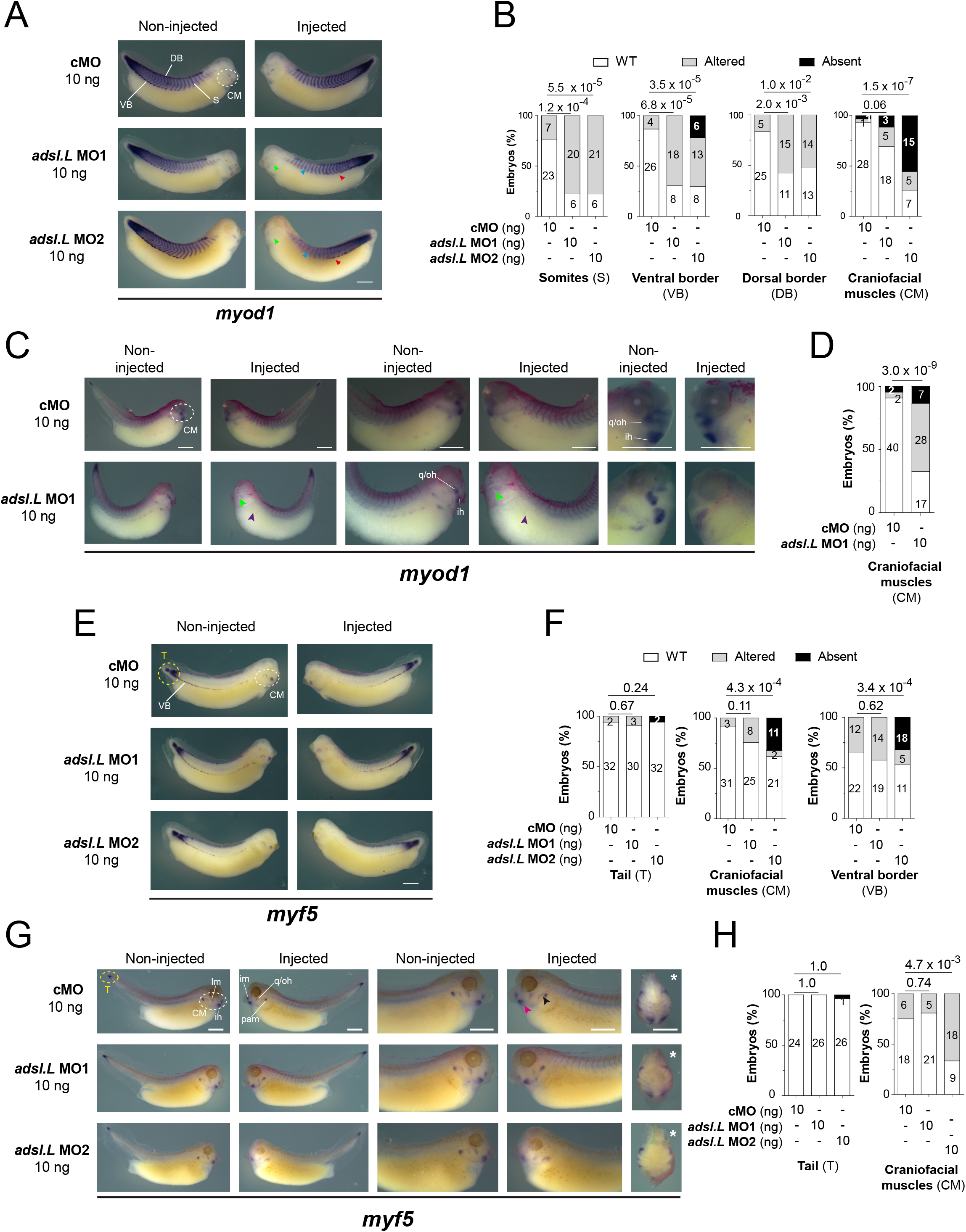
Knock-down of *adsl.L* gene causes an impaired expression of *myod1* and *myf5* at late tailbud and tadpole embryonic stages leading to somites and craniofacial muscle formation defects. (**A-D**) Representative images of the *adsl.L*-dependent altered expression of *myod1* (**A**) and *myf5* (**C**) domains in either somite (S), dorsal somite border (DB), ventral somite border (VB) or craniofacial muscles (CM), as revealed by *in situ* hybridization in late tailbud embryos. (**E**, **G**) Representative images of the *adsl.L*-dependent altered expression of *myod1* (**E**) and *myf5* (**G**) domains in the unsegmented mesoderm in the most posterior tail (T), somites (S), ventral somite border (VB) and craniofacial muscles (CM) in tadpole embryos. (**B, D, F, H**) Quantification and statistics of the *myod1* and *myf5* expression phenotypes are presented in (**A**), (**C**), (**E**) and (**G**), respectively. Craniofacial muscles: ih, interhyoïedus anlage; im, intermandibularis anlage; lm, levatores mandibulae anlage; pam, pharyngeal archmuscle anlagen; q/oh, common orbitohyoïdeus and quadrato-hyoangularis precursors. Injected side is indicated by asterisks. Bars: 0.5 mm. Numbers above the bars of histograms correspond to p-values.

**Figure 6.**
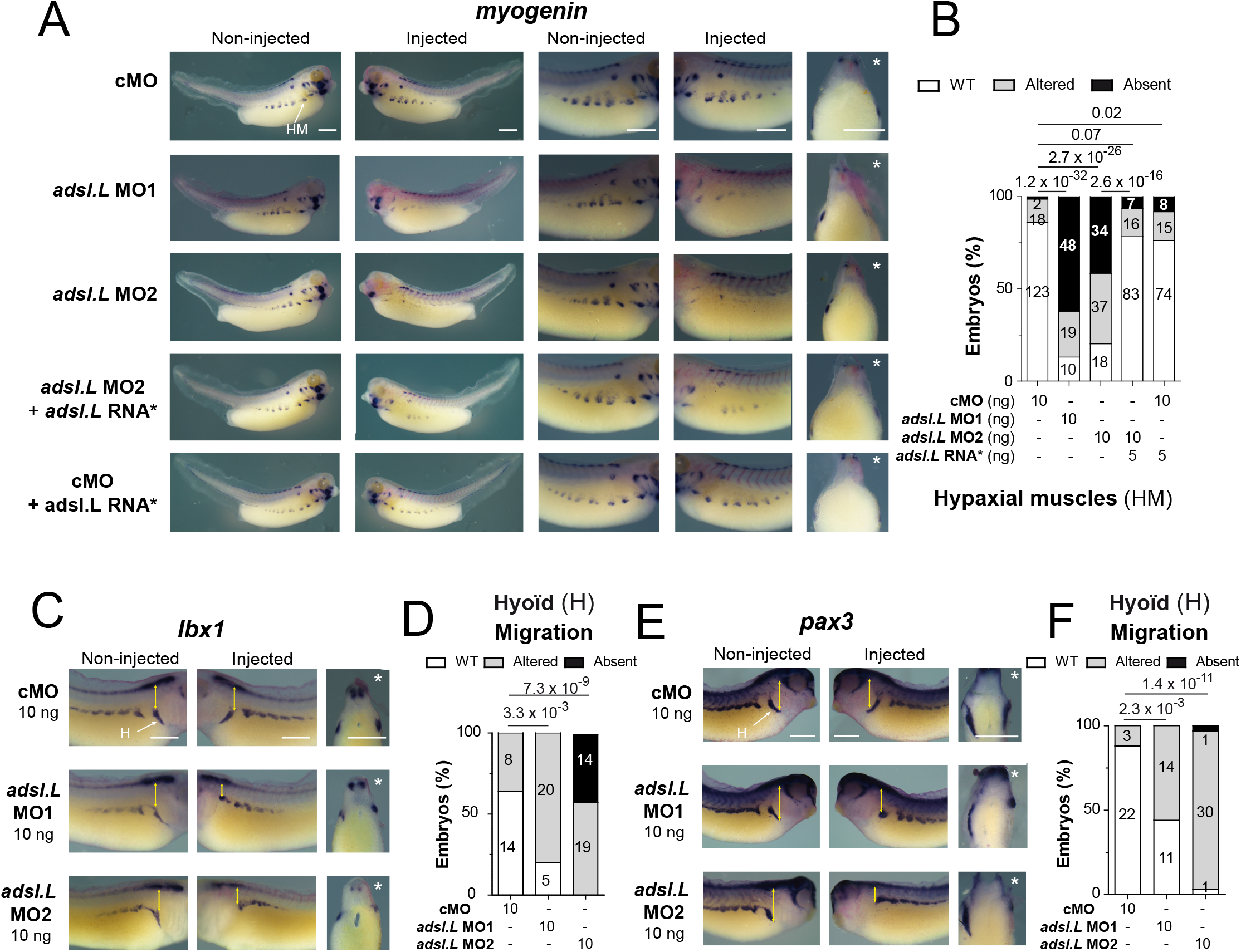
The *adsl.L* gene is required for hypaxial muscle migration. (**A-B**) The hypaxial muscle defect associated with *adsl.L* knock-down is rescued by the MO non-targeted *adsl.L* RNA*. Representative images (**A**), quantifications (number in bars) and statistics (**B**) of the *adsl.L*-dependent alteration of *myogenin* gene expression monitored by *in situ* hybridization. (**C-F**) Migration of myoblasts in the hyoid region is found severely affected in *adsl.L* morphants as shown by *lbx1* (**C-D**) and *pax3* genes expression pattern alteration (**E-F**). Injected side is indicated by asterisks. Bars: 0.5 mm. HM and H stand for hypaxial muscles and hyoid region, respectively Numbers above the bars of histograms correspond to p-values.

### 3.5 The adsl.L gene is required for hypaxial muscle formation and migration

In addition to the somite and craniofacial muscle phenotypes, severe hypaxial muscle alterations were observed in *adsl.L* morphants (Figure 3 and Figure S9). To further characterize these defects, the expression of *myogenin* was analyzed by *in situ* hybridization at the late tadpole stage. *Adsl.L* knock-down resulted in either a strong decrease or even an absence of *myog* expression in hypaxial muscle cells, while no significant alteration was observed in the embryos injected with the control morpholino (cMO) in combination or not with the *adsl.L* mutated RNA* (Figure 6A-B). This muscle phenotype was rescued by the injection of the *adsl.L* mutated RNA* in *adsl.L* MO2 morphants. These *myogenin* expression alterations were confirmed as *myod1* expression is strongly reduced or absent in hypaxial muscles in *adsl.L* morphants (purple arrowheads, Figure 5C and Figure S15). These results confirm the role of the *adsl.L* gene in the expression of myogenic regulatory factors required for correct hypaxial muscle development. In addition, hypaxial muscles were also found to be highly reduced or absent as revealed by 12-101 immunolabelling in *ppat.L/ppat.S* and *hprt1.L* morphants (pink arrowheads, Figure S10-S11).

To better understand the deleterious effect of *adsl.L* knock-down on hypaxial muscles, expression of the *pax3* and *lbx1* genes encoding transcription factors were analyzed at tadpole stages. *Lbx1* and *pax3* are both expressed in hypaxial body wall undifferentiated cells that form the front of hypaxial myoblast migration [37,48]. Although the injection of *adsl.L* MO, especially *adsl.L* MO2, induces a slight reduction of *lbx1* expression in the hypaxial muscle precursors, the major observed phenotype is a migration alteration of these *lbx1* positive cells (Figure 6C-D). Indeed, muscle progenitors are closer to the ventral border of the somites. This alteration is more pronounced in the most anterior region (trunk I), in which progenitors migrate around the hyoid region (see yellow double arrows in Figure 6C). Similar alterations were observed for *pax3* positive hypaxial body wall progenitors, characterized by a severe alteration of hypaxial progenitor migration, while *pax3* expression is less or not affected in the hypaxial myotome (Figure 6E-F). Altogether, these results show that purine biosynthesis genes are essential during hypaxial muscle formation in *X. laevis*.

## 4. Discussion

### 4.1 X. laevis as a vertebrate model to study purine deficiency related pathologies

In this study, we functionally characterized for the first time the main members of the purine biosynthesis and salvage pathways in the vertebrate *X. laevis*. We also established the comparative map of their embryonic spatiotemporal expression and showed that the main common expression of purine genes is in the neuro-muscular tissue and its precursors. This expression profile can be directly linked to the neuro-muscular symptoms usually observed in most of the nearly thirty identified purine-associated pathologies [3–7]. Furthermore, we investigated the embryonic roles of three purine biosynthesis enzymes during vertebrate muscle development, focusing on the Adsl.L enzyme. As *ADSL* deficiency in humans is associated with a residual enzymatic activity (reviewed in [6,49]), we believe that our experimental strategy based on the knockdown with morpholino-oligonucleotides is more appropriate in identifying the molecular dysfunctions found in patients rather than strategies based on knock-out experiments. Altogether, our data shows that *X. laevis* is a relevant model in studying the molecular bases of the developmental alterations associated with purine deficiencies.

### 4.2 Role of purine pathway genes during X. laevis development

Maternal and zygotic expression are key indicators of potential pathway involvement during vertebrate development. We show that the early stages of *X. laevis* embryo development is dependent on both purine pathways, as all the genes studied, except for *adss1.L* and *adss1.S,* are maternal genes and are expressed from gastrula stages. However, from the early organogenesis phase (stage 22), all the 17 tested genes are expressed. We have previously shown that the unique reserve of purine in embryos, hypoxanthine, is depleted at stage 22 [46], so this embryonic stage is placed as a key turning point of purine source for the embryo. From this stage onwards and until stage 45 (stage where embryos start to feed), purines can only be synthesized *de novo* or recycled within the embryo, and zygotic expression of purine biosynthesis genes becomes then crucial.

Furthermore, the genes of the purine *de novo* and salvage pathways share simultaneous embryonic expression and mostly in the same tissues, strongly suggesting that neither pathway is preferred for purine supply during *X. laevis* development. Our functional experiments also demonstrate that both pathways are involved in development since the muscle alterations observed in *adsl.L* morphants were also observed in *ppat.L/ppat.S* and *hprt1.L* morphants. This is different from what has been obtained in *C. elegans* in which muscle phenotypes associated with *ADSL* deficiency are rather dependent on the purine recycling pathway [27].

In addition, we have shown that *adsl.L* knockdown alters myogenesis and is associated with strong alterations in anterior somite formation and subsequent malformation of hypaxial and craniofacial muscles. Embryonic neuro-muscular defects have previously been described in other animal models [26,27], with a strong alteration in muscle integrity observed in *C. elegans adsl* knockout mutants. These studies raised the question of whether the observed phenotypes were the result of a decrease in one or more end products of the purine biosynthetic pathways, or that of an accumulation of Adsl.L substrates or their derivatives. Indeed, the neurodevelopmental alterations observed in zebrafish *adsl* mutants were associated with the accumulation of SAICAr, the dephosphorylated form of Adsl substrate in the *de novo* pathway (SAICAR, or SZMP) and thus these neuronal alterations were rescued by inhibition of this pathway [26]. In *X. laevis*, as the alterations in myogenesis are very similar in the *adsl.L*, *ppat.L*/*ppat.S* and *hprt1.L* morphants, it seems very unlikely that the muscle phenotypes are related to an accumulation of Adsl.L substrates or their derivatives (Figure 1A). A metabolic rescue of the phenotypes is, therefore, an appropriate strategy to determine whether the muscle phenotypes could be related to a decrease in Adsl.L products and/or downstream metabolites. Hypoxanthine, the main source of purine in *X. laevis* embryo [46], would have been the best candidate metabolite for the MO rescue but it cannot be tested here since its metabolization requires the enzymatic activity of Adsl.L (Figure 1A). Rescue experiments were therefore carried out by adding adenine to the Adsl.L morphant culture medium in concentrations at which this purine precursor was soluble in the medium (50 µM) but without any success. We cannot rule out that this may be due to the incapacity of this purine to cross the vitelline membrane. However, rescue experiments by adenine injection into the blastocoel or archenteron, as previously performed for other purine [32,50], is not possible because of its low solubility in aqueous solutions (< few µM).

Overall, these results strongly suggest that the observed muscle phenotypes are associated with purine deficiency, regardless of the mode of biosynthesis of these purines, since disruption of the *de novo* pathway (*ppat.L*/*ppat.S* MO), the recycling pathway (*hprt1.L* MO), or both (*adsl.L* MO) results in similar alterations in myogenesis. However, these alterations may explain the muscle symptoms such as axial hypotonia, peripheral hypotonia and muscle wasting observed in patients [13,16,17].

### 4.3 Muscle defects associated with purine pathways dysfunction are related to an altered expression of MRF genes

Our study provides new *in vivo* evidence for the identification of altered molecular mechanisms during purine deficiency. Indeed, we show that functional purine pathways are required for the expression of the myogenic regulator factors MyoD1, Myf5 and Myogenin, which are by far the master regulatory transcription factors of the embryonic myogenic program including myotome and somite formation, hypaxial and craniofacial myogenesis [36,47]. The *adsl.L*-specific alteration of *myod1* and *myf5* expression in the ventral border of the dermomyotome could, therefore, be at the origin of the deleterious effects observed in morphants on hypaxial progenitors and differentiated hypaxial muscle cells. A strong decrease in MRF expression was observed in *adsl.L* morphants from the earliest stages of muscle formation in neurula embryos. As MRF genes regulate the expression of other MRFs and/or their own expression, it is possible that one (or more) key purine derivative(s) could regulate the transcription expression and/or the function of one of the MRF. Interestingly, the opposite phenotypes were obtained on *myod1* expression in *adsl.L* morphants and in embryos overexpressing this gene, placing *adsl.L* as a key regulator of *myod1* expression during development. Since MyoD1 is a potent inducer of muscle differentiation and is required for *myf5* expression in early *X. laevis* development, correct anterior somite segmentation [51] and for correct myotome size [35], it is, therefore, possible that the muscle defects in *adsl.L* morphants are mainly due to a deregulation of *myod1* expression, thus placing this gene, and the purine biosynthetic pathways, at the top of the myogenic gene cascade. An interesting future point of investigation would be to test whether Myod1 expression is able to rescue adsl morphant muscle phenotypes.

So far, we have not found a direct nor an indirect link between purines and the transcriptional regulation of MRFs. However, regulation of transcription factor activity by direct binding of purine metabolites has already been demonstrated in different organisms [22,52]. In yeast, we have shown that the direct binding of purine metabolites (SZMP and ZMP, Figure 1) on Pho4 and Pho2 transcription factors was responsible for their activation [22]. These two metabolites of the *de novo* purine pathway are certainly not those implicated here as similar muscle phenotypes were obtained in *adsl.L*, *ppat.L*/*ppat.S* and *hprt1.L* morphants, but we can speculate that another purine derivative could similarly be involved in modulating the transcriptional expression of MRFs during *X. laevis* myogenesis. Moreover, regulation of *myogenin* gene expression by metabolites (CMP and UMP) has recently been observed in the mouse C2C12 cell line [53], showing that MRF expression can be modulated by nucleotides. To our knowledge, a similar direct regulation of MRFs by purines has not been described to date. If it exists, as suggested by our data, then the mechanism by which it acts remains to be established.

It may be possible that the effect of purines on MRFs expression is indirect via upstream cascades, responding to purines, and regulating the expression of these genes. Indeed, many upstream signals, for example, Wnt and FGF, regulate the expression of MRFs in myogenesis (reviewed in [54]). These pathways involve the activation of protein kinases of the myogenic kinome whose activity is highly dependent on intracellular pools of purine nucleotides [55]. In particular, the activity of protein kinase A, initiated by the activation of heteromeric G proteins and adenylate cyclase through the Wnt pathway, is required for the expression of *myod1* and *myf5* and the formation of myoblasts. In addition, extracellular triphosphate purine nucleotides are ligands of purinergic receptors which are involved in myoblast proliferation and differentiation [56,57]. We have recently shown that the ATP-dependent P2X5 receptor subunit is specifically expressed in *X. laevis* somites, which is in agreement with its expression profile described in other animal models [58,59]. We may hypothesize that *adsl.L* knock-down may reduce the availability of extracellular purines, leading to a possible alteration in P2X5 activation and myogenesis impairment [59]. This hypothesis will be tested in the next future in the laboratory. We had previously shown that purinergic signaling controls vertebrate eye development by acting on the expression of the PSED (*pax*/*six*/*eya*/*dach*) network genes [29]. As this PSED gene network regulates MRF expression [60], it may be also possible that a similar mechanism is involved in *adsl.L* morphants. Interestingly, our observed muscle phenotypes are highly similar to those induced by retinoic acid receptor β2 (RARβ2) loss of function [61]. Retinoic acid pathway, through the activation of RAR, regulates myogenic differentiation and MRF gene expression [62]. It has been reported that retinoic acid affects synthesis of PRPP in human erythrocytes in Psoriasis [63] and RA deficiency interferes with the expression of genes involved in the purine metabolism pathway *in vivo* [64]. These data along with ours suggest a possible link between purine biosynthetic pathway, retinoic acid signaling pathway and MRF gene expression, worth of further studies in the future.

In conclusion, the establishment of this new animal model allowed us to demonstrate the critical functions of the purine biosynthetic pathways during vertebrate embryogenesis. Indeed, our results provide evidence for the roles for purine pathway genes in myogenesis through the regulation of MRF gene expression during embryogenesis. Although the first median and lateral myogenesis disappeared during evolution, the second myogenic wave is conserved [47]. We can, therefore, speculate that the involvement of the *de novo* and salvage purine pathways during this myogenic wave is conserved in mammals and that purines may, therefore, be key regulators in the formation of hypaxial myogenic cells, the progenitors of the abdominal, spinal, and limb muscles, those muscles which are affected in patients with purine-associated pathologies [4,5,7,49]. Although, in this work we have mostly focused on muscle alterations, it provides the basis for further functional studies to identify the molecular mechanisms involved in the development of neuromuscular tissues, those tissues in which the alterations underlie the major deleterious symptoms of patients with purine deficiency.

## Author contributions

Conceptualization, B.P. and K.M.; methodology, B.P., C.S-M, E.H., K.M. and M.D..; Formal analysis, B.P., K.M. and M.D.; Investigation, B.P., C.S-M, E.H., K.M. and M.D..; Resources, B.D-F, B.P., C.S-M, E. B-G., E.H., K.M. and M.D.; Writing-original draft preparation, B.P., K.M.; Supervision, B.P. and K.M. All authors have read and agreed to the published version of the manuscript.

## Acknowledgments

The author thanks C. Blanchard, A. Martinez and A. Tocco for their technical help, D. Patterson for generous gift of the pADSLpCR3.1 plasmid used to clone human *ADSL*-pCS2+ plasmid for the rescue experiments, C. Chanoine for generous gift of the *myogenin* plasmid used for *in situ* hybridization probes, J.E. Gomes for critical reading of the manuscript, and K. Chiimba and S. Sanghera for English proofreading.

## References

1. Daignan-Fornier, B.; Pinson, B. Yeast to Study Human Purine Metabolism Diseases. Cells 2019, 8, E67, doi:10.3390/cells8010067.

2. Massé, K.; Dale, N. Purines as Potential Morphogens during Embryonic Development. Purinergic Signal. 2012, 8, 503–521, doi:10.1007/s11302-012-9290-y.

3. Simmonds, H.A.; Duley, J.A.; Fairbanks, L.D.; McBride, M.B. When to Investigate for Purine and Pyrimidine Disorders. Introduction and Review of Clinical and Laboratory Indications. J. Inherit. Metab. Dis. 1997, 20, 214–226, doi:10.1023/a:1005308923168.

4. Balasubramaniam, S.; Duley, J.A.; Christodoulou, J. Inborn Errors of Purine Metabolism: Clinical Update and Therapies. J. Inherit. Metab. Dis. 2014, 37, 669–686, doi:10.1007/s10545-014-9731-6.

5. Jurecka, A. Inborn Errors of Purine and Pyrimidine Metabolism. J. Inherit. Metab. Dis. 2009, 32, 247–263, doi:10.1007/s10545-009-1094-z.

6. Nyhan, W.L. Disorders of Purine and Pyrimidine Metabolism. Mol. Genet. Metab. 2005, 86, 25–33, doi:10.1016/j.ymgme.2005.07.027.

7. van den Berghe, G.; Vincent, M.-F.; Marie, S. Disorders of Purine and Pyrimidine Metabolism. In Inborn Metabolic Diseases: Diagnosis and Treatment; Fernandes, J., Saudubray, J.-M., van den Berghe, G., Walter, J.H., Eds.; Springer: Berlin, Heidelberg, 2006; pp. 433–449 ISBN 978-3-540-28785-8.

8. Jaeken, J.; Van den Berghe, G. An Infantile Autistic Syndrome Characterised by the Presence of Succinylpurines in Body Fluids. Lancet Lond. Engl. 1984, 2, 1058–1061.

9. Stone, R.L.; Aimi, J.; Barshop, B.A.; Jaeken, J.; Van den Berghe, G.; Zalkin, H.; Dixon, J.E. A Mutation in Adenylosuccinate Lyase Associated with Mental Retardation and Autistic Features. Nat. Genet. 1992, 1, 59–63, doi:10.1038/ng0492-59.

10. Mao, X.; Li, K.; Tang, B.; Luo, Y.; Ding, D.; Zhao, Y.; Wang, C.; Zhou, X.; Liu, Z.; Zhang, Y.;, et al. Novel Mutations in ADSL for Adenylosuccinate Lyase Deficiency Identified by the Combination of Trio-WES and Constantly Updated Guidelines. Sci. Rep. 2017, 7, 1625, doi:10.1038/s41598-017-01637-z.

11. Van den Bergh, F.; Vincent, M.F.; Jaeken, J.; Van den Berghe, G. Residual Adenylosuccinase Activities in Fibroblasts of Adenylosuccinase-Deficient Children: Parallel Deficiency with Adenylosuccinate and Succinyl-AICAR in Profoundly Retarded Patients and Non-Parallel Deficiency in a Mildly Retarded Girl. J. Inherit. Metab. Dis. 1993, 16, 415–424, doi:10.1007/BF00710291.

12. Kmoch, S.; Hartmannová, H.; Stibůrková, B.; Krijt, J.; Zikánová, M.; Sebesta, I. Human Adenylosuccinate Lyase (ADSL), Cloning and Characterization of Full-Length CDNA and Its Isoform, Gene Structure and Molecular Basis for ADSL Deficiency in Six Patients. Hum. Mol. Genet. 2000, 9, 1501–1513, doi:10.1093/hmg/9.10.1501.

13. Mouchegh, K.; Zikánová, M.; Hoffmann, G.F.; Kretzschmar, B.; Kühn, T.; Mildenberger, E.; Stoltenburg-Didinger, G.; Krijt, J.; Dvoráková, L.; Honzík, T.;, et al. Lethal Fetal and Early Neonatal Presentation of Adenylosuccinate Lyase Deficiency: Observation of 6 Patients in 4 Families. J. Pediatr. 2007, 150, 57–61.e2, doi:10.1016/j.jpeds.2006.09.027.

14. Jurecka, A.; Zikanova, M.; Tylki-Szymanska, A.; Krijt, J.; Bogdanska, A.; Gradowska, W.; Mullerova, K.; Sykut-Cegielska, J.; Kmoch, S.; Pronicka, E. Clinical, Biochemical and Molecular Findings in Seven Polish Patients with Adenylosuccinate Lyase Deficiency. Mol. Genet. Metab. 2008, 94, 435–442, doi:10.1016/j.ymgme.2008.04.013.

15. Zikanova, M.; Skopova, V.; Hnizda, A.; Krijt, J.; Kmoch, S. Biochemical and Structural Analysis of 14 Mutant Adsl Enzyme Complexes and Correlation to Phenotypic Heterogeneity of Adenylosuccinate Lyase Deficiency. Hum. Mutat. 2010, 31, 445–455, doi:10.1002/humu.21212.

16. Spiegel, E.K.; Colman, R.F.; Patterson, D. Adenylosuccinate Lyase Deficiency. Mol. Genet. Metab. 2006, 89, 19–31, doi:10.1016/j.ymgme.2006.04.018.

17. Gitiaux, C.; Ceballos-Picot, I.; Marie, S.; Valayannopoulos, V.; Rio, M.; Verrieres, S.; Benoist, J.F.; Vincent, M.F.; Desguerre, I.; Bahi-Buisson, N. Misleading Behavioural Phenotype with Adenylosuccinate Lyase Deficiency. Eur. J. Hum. Genet. EJHG 2009, 17, 133–136, doi:10.1038/ejhg.2008.174.

18. Jurecka, A.; Zikanova, M.; Kmoch, S.; Tylki-Szymańska, A. Adenylosuccinate Lyase Deficiency. J. Inherit. Metab. Dis. 2015, 38, 231–242, doi:10.1007/s10545-014-9755-y.

19. Marie, S.; Race, V.; Vincent, M.F.; Van den Berghe, G. Adenylosuccinate Lyase Deficiency: From the Clinics to Molecular Biology. Adv. Exp. Med. Biol. 2000, 486, 79– 82, doi:10.1007/0-306-46843-3_15.

20. Race, V.; Marie, S.; Vincent, M.F.; Van den Berghe, G. Clinical, Biochemical and Molecular Genetic Correlations in Adenylosuccinate Lyase Deficiency. Hum. Mol. Genet. 2000, 9, 2159–2165, doi:10.1093/hmg/9.14.2159.

21. Mastrogiorgio, G.; Macchiaiolo, M.; Buonuomo, P.S.; Bellacchio, E.; Bordi, M.; Vecchio, D.; Brown, K.P.; Watson, N.K.; Contardi, B.; Cecconi, F.;, et al. Clinical and Molecular Characterization of Patients with Adenylosuccinate Lyase Deficiency. Orphanet J. Rare Dis. 2021, 16, 112, doi:10.1186/s13023-021-01731-6.

22. Pinson, B.; Vaur, S.; Sagot, I.; Coulpier, F.; Lemoine, S.; Daignan-Fornier, B. Metabolic Intermediates Selectively Stimulate Transcription Factor Interaction and Modulate Phosphate and Purine Pathways. Genes Dev. 2009, 23, 1399–1407, doi:10.1101/gad.521809.

23. Keller, K.E.; Tan, I.S.; Lee, Y.-S. SAICAR Stimulates Pyruvate Kinase Isoform M2 and Promotes Cancer Cell Survival in Glucose-Limited Conditions. Science 2012, 338, 1069–1072, doi:10.1126/science.1224409.

24. Gooding, J.R.; Jensen, M.V.; Dai, X.; Wenner, B.R.; Lu, D.; Arumugam, R.; Ferdaoussi, M.; MacDonald, P.E.; Newgard, C.B. Adenylosuccinate Is an Insulin Secretagogue Derived from Glucose-Induced Purine Metabolism. Cell Rep. 2015, 13, 157–167, doi:10.1016/j.celrep.2015.08.072.

25. Mazzarino, R.C.; Baresova, V.; Zikánová, M.; Duval, N.; Wilkinson, T.G.; Patterson, D.; Vacano, G.N. The CRISPR-Cas9 CrADSL HeLa Transcriptome: A First Step in Establishing a Model for ADSL Deficiency and SAICAR Accumulation. Mol. Genet. Metab. Rep. 2019, 21, 100512, doi:10.1016/j.ymgmr.2019.100512.

26. Dutto, I.; Gerhards, J.; Herrera, A.; Souckova, O.; Škopová, V.; Smak, J.A.; Junza, A.; Yanes, O.; Boeckx, C.; Burkhalter, M.D.;, et al. Pathway-Specific Effects of ADSL Deficiency on Neurodevelopment. eLife 2022, 11, e70518, doi:10.7554/eLife.70518.

27. Marsac, R.; Pinson, B.; Saint-Marc, C.; Olmedo, M.; Artal-Sanz, M.; Daignan-Fornier, B.; Gomes, J.-E. Purine Homeostasis Is Necessary for Developmental Timing, Germline Maintenance and Muscle Integrity in Caenorhabditis Elegans. Genetics 2019, 211, 1297–1313, doi:10.1534/genetics.118.301062.

28. Sater, A.K.; Moody, S.A. Using Xenopus to Understand Human Disease and Developmental Disorders. Genes. N. Y. N 2000 2017, 55, doi:10.1002/dvg.22997.

29. Blum, M.; Ott, T. Xenopus: An Undervalued Model Organism to Study and Model Human Genetic Disease. Cells Tissues Organs 2018, 205, 303–313, doi:10.1159/000490898.

30. Garí, E.; Piedrafita, L.; Aldea, M.; Herrero, E. A Set of Vectors with a Tetracycline-Regulatable Promoter System for Modulated Gene Expression in Saccharomyces Cerevisiae. Yeast Chichester Engl. 1997, 13, 837–848, doi:10.1002/(SICI)1097-0061(199707)13:9&lt;837::AID-YEA145>3.0.CO;2-T.

31. Nieuwkoop, P.D.; Faber, J. Normal Table of Xenopus Laevis (Daudin): A Systematical and Chronological Survey of the Development from the Fertilized Egg Till the End of Metamorphosis; Garland Pub., 1994; ISBN 978-0-8153-1896-5.

32. Massé, K.; Bhamra, S.; Eason, R.; Dale, N.; Jones, E.A. Purine-Mediated Signalling Triggers Eye Development. Nature 2007, 449, 1058–1062, doi:10.1038/nature06189.

33. Blanchard, C.; Boué-Grabot, E.; Massé, K. Comparative Embryonic Spatio-Temporal Expression Profile Map of the Xenopus P2X Receptor Family. Front. Cell. Neurosci. 2019, 13, 340, doi:10.3389/fncel.2019.00340.

34. Massé, K.; Eason, R.; Bhamra, S.; Dale, N.; Jones, E.A. Comparative Genomic and Expression Analysis of the Conserved NTPDase Gene Family in Xenopus. Genomics 2006, 87, 366–381, doi:10.1016/j.ygeno.2005.11.003.

35. Della Gaspera, B.; Armand, A.-S.; Lecolle, S.; Charbonnier, F.; Chanoine, C. Mef2d Acts Upstream of Muscle Identity Genes and Couples Lateral Myogenesis to Dermomyotome Formation in Xenopus Laevis. PloS One 2012, 7, e52359, doi:10.1371/journal.pone.0052359.

36. Della Gaspera, B.; Armand, A.-S.; Sequeira, I.; Chesneau, A.; Mazabraud, A.; Lécolle, S.; Charbonnier, F.; Chanoine, C. Myogenic Waves and Myogenic Programs during Xenopus Embryonic Myogenesis. Dev. Dyn. Off. Publ. Am. Assoc. Anat. 2012, 241, 995–1007, doi:10.1002/dvdy.23780.

37. Martin, B.L.; Harland, R.M. A Novel Role for Lbx1 in Xenopus Hypaxial Myogenesis. Dev. Camb. Engl. 2006, 133, 195–208, doi:10.1242/dev.02183.

38. Kintner, C.R.; Brockes, J.P. Monoclonal Antibodies Identify Blastemal Cells Derived from Dedifferentiating Limb Regeneration. Nature 1984, 308, 67–69, doi:10.1038/308067a0.

39. Massé, K.; Bhamra, S.; Allsop, G.; Dale, N.; Jones, E.A. Ectophosphodiesterase/Nucleotide Phosphohydrolase (Enpp) Nucleotidases: Cloning, Conservation and Developmental Restriction. Int. J. Dev. Biol. 2010, 54, 181–193, doi:10.1387/ijdb.092879km.

40. Bassez, T.; Paris, J.; Omilli, F.; Dorel, C.; Osborne, H.B. Post-Transcriptional Regulation of Ornithine Decarboxylase in Xenopus Laevis Oocytes. Dev. Camb. Engl. 1990, 110, 955–962.

41. Bowes, J.B.; Snyder, K.A.; Segerdell, E.; Gibb, R.; Jarabek, C.; Noumen, E.; Pollet, N.; Vize, P.D. Xenbase: A Xenopus Biology and Genomics Resource. Nucleic Acids Res. 2008, 36, D761–767, doi:10.1093/nar/gkm826.

42. Karpinka, J.B.; Fortriede, J.D.; Burns, K.A.; James-Zorn, C.; Ponferrada, V.G.; Lee, J.; Karimi, K.; Zorn, A.M.; Vize, P.D. Xenbase, the Xenopus Model Organism Database; New Virtualized System, Data Types and Genomes. Nucleic Acids Res. 2015, 43, D756–763, doi:10.1093/nar/gku956.

43. Session, A.M.; Uno, Y.; Kwon, T.; Chapman, J.A.; Toyoda, A.; Takahashi, S.; Fukui, A.; Hikosaka, A.; Suzuki, A.; Kondo, M.;, et al. Genome Evolution in the Allotetraploid Frog Xenopus Laevis. Nature 2016, 538, 336–343, doi:10.1038/nature19840.

44. Ljungdahl, P.O.; Daignan-Fornier, B. Regulation of Amino Acid, Nucleotide, and Phosphate Metabolism in Saccharomyces Cerevisiae. Genetics 2012, 190, 885–929, doi:10.1534/genetics.111.133306.

45. Kragtorp, K.A.; Miller, J.R. Integrin Alpha5 Is Required for Somite Rotation and Boundary Formation in Xenopus. Dev. Dyn. Off. Publ. Am. Assoc. Anat. 2007, 236, 2713–2720, doi:10.1002/dvdy.21280.

46. Tocco, A.; Pinson, B.; Thiébaud, P.; Thézé, N.; Massé, K. Comparative Genomic and Expression Analysis of the Adenosine Signaling Pathway Members in Xenopus. Purinergic Signal. 2015, 11, 59–77, doi:10.1007/s11302-014-9431-6.

47. Della Gaspera, B.; Weill, L.; Chanoine, C. Evolution of Somite Compartmentalization: A View From Xenopus. Front. Cell Dev. Biol. 2021, 9, 790847, doi:10.3389/fcell.2021.790847.

48. Martin, B.L.; Harland, R.M. Hypaxial Muscle Migration during Primary Myogenesis in Xenopus Laevis. Dev. Biol. 2001, 239, 270–280, doi:10.1006/dbio.2001.0434.

49. Kelley, R.E.; Andersson, H.C. Disorders of Purines and Pyrimidines. Handb. Clin. Neurol. 2014, 120, 827–838, doi:10.1016/B978-0-7020-4087-0.00055-3.

50. Iijima, R.; Kunieda, T.; Yamaguchi, S.; Kamigaki, H.; Fujii-Taira, I.; Sekimizu, K.; Kubo, T.; Natori, S.; Homma, K.J. The Extracellular Adenosine Deaminase Growth Factor, ADGF/CECR1, Plays a Role in Xenopus Embryogenesis via the Adenosine/P1 Receptor. J. Biol. Chem. 2008, 283, 2255–2264, doi:10.1074/jbc.M709279200.

51. Maguire, R.J.; Isaacs, H.V.; Pownall, M.E. Early Transcriptional Targets of MyoD Link Myogenesis and Somitogenesis. Dev. Biol. 2012, 371, 256–268, doi:10.1016/j.ydbio.2012.08.027.

52. Kim, P.B.; Nelson, J.W.; Breaker, R.R. An Ancient Riboswitch Class in Bacteria Regulates Purine Biosynthesis and One-Carbon Metabolism. Mol. Cell 2015, 57, 317– 328, doi:10.1016/j.molcel.2015.01.001.

53. Nakagawara, K.; Takeuchi, C.; Ishige, K. 5’-CMP and 5’-UMP Promote Myogenic Differentiation and Mitochondrial Biogenesis by Activating Myogenin and PGC-1α in a Mouse Myoblast C2C12 Cell Line. Biochem. Biophys. Rep. 2022, 31, 101309, doi:10.1016/j.bbrep.2022.101309.

54. Sabillo, A.; Ramirez, J.; Domingo, C.R. Making Muscle: Morphogenetic Movements and Molecular Mechanisms of Myogenesis in Xenopus Laevis. Semin. Cell Dev. Biol. 2016, 51, 80–91, doi:10.1016/j.semcdb.2016.02.006.

55. Knight, J.D.; Kothary, R. The Myogenic Kinome: Protein Kinases Critical to Mammalian Skeletal Myogenesis. Skelet. Muscle 2011, 1, 29, doi:10.1186/2044-5040-1-29.

56. Burnstock, G.; Arnett, T.R.; Orriss, I.R. Purinergic Signalling in the Musculoskeletal System. Purinergic Signal. 2013, 9, 541–572, doi:10.1007/s11302-013-9381-4.

57. Pietrangelo, T.; Guarnieri, S.; Fulle, S.; Fanò, G.; Mariggiò, M.A. Signal Transduction Events Induced by Extracellular Guanosine 5’ Triphosphate in Excitable Cells. Purinergic Signal. 2006, 2, 633–636, doi:10.1007/s11302-006-9021-3.

58. Meyer, M.P.; Gröschel-Stewart, U.; Robson, T.; Burnstock, G. Expression of Two ATP-Gated Ion Channels, P2X5 and P2X6, in Developing Chick Skeletal Muscle. Dev. Dyn. Off. Publ. Am. Assoc. Anat. 1999, 216, 442–449, doi:10.1002/(SICI)1097-0177(199912)216:4/5&lt;442::AID-DVDY12>3.0.CO;2-Z.

59. Ryten, M.; Dunn, P.M.; Neary, J.T.; Burnstock, G. ATP Regulates the Differentiation of Mammalian Skeletal Muscle by Activation of a P2X5 Receptor on Satellite Cells. J. Cell Biol. 2002, 158, 345–355, doi:10.1083/jcb.200202025.

60. Maire, P.; Dos Santos, M.; Madani, R.; Sakakibara, I.; Viaut, C.; Wurmser, M. Myogenesis Control by SIX Transcriptional Complexes. Semin. Cell Dev. Biol. 2020, 104, 51–64, doi:10.1016/j.semcdb.2020.03.003.

61. Janesick, A.; Tang, W.; Nguyen, T.T.L.; Blumberg, B. RARβ2 Is Required for Vertebrate Somitogenesis. Dev. Camb. Engl. 2017, 144, 1997–2008, doi:10.1242/dev.144345.

62. Chen, J.; Li, Q. Implication of Retinoic Acid Receptor Selective Signaling in Myogenic Differentiation. Sci. Rep. 2016, 6, 18856, doi:10.1038/srep18856.

63. Baló-Banga, J.M.; Leibinger, J.; Molnár, L.; Király, K. The Effect of Retinoic Acid on the Synthesis of Phosphoribosylpyrophosphate in Human Erythrocytes in Psoriasis. A Preliminary Note. Dermatologica 1978, 157 Suppl 1, 45–51, doi:10.1159/000250884.

64. Paschaki, M.; Schneider, C.; Rhinn, M.; Thibault-Carpentier, C.; Dembélé, D.; Niederreither, K.; Dollé, P. Transcriptomic Analysis of Murine Embryos Lacking Endogenous Retinoic Acid Signaling. PloS One 2013, 8, e62274, doi:10.1371/journal.pone.0062274.

