## Supplemental material for "Purine biosynthesis pathways are required for myogenesis in *Xenopus laevis*"

### Supplementary Figures

**Figure S1**. Conservation of purine biosynthesis pathways between the yeast *Saccharomyces cerevisiae* and *Xenopus laevis* p. 2
**Figure S2**. Sequence comparison between the *H. sapiens* (*Hs*) and *X. laevis* (*Xl*) adenylosuccinate lyase enzymes p. 3
**Figure S3.** Validation of the morpholinos translation interference by *in vitro* translation p. 4
**Figure S4.** Functional complementation of yeast purine *de novo* pathway mutants by the *X. laevis* ortholog genes p. 5
**Figure S5.** Functional complementation of yeast purine salvage knocked-out mutants by the *X. laevis* ortholog genes p. 6
**Figure S6.** Temporal expression profiles of purine pathway genes during embryogenesis p. 7
**Figure S7.** The expression profile was determined for each gene by *in situ* hybridization (ISH) using antisense probes p. 8
**Figure S8.** *In situ* hybridization using control sense probes p. 9
**Figure S9.** Severity of *adsl.L* knock-down-associated phenotypes is morpholino dose-dependent p. 10
**Figure S10.** The *ppat.L* and *ppat.S* genes are required for somites and hypaxial muscle formation in *X. laevis* p. 11
**Figure S11.** The *hprt1.L* gene is required for somitogenesis and hypaxial muscle formation in *X. laevis* p. 12
**Figure S12.** Effect of *H. sapiens ADSL* and *X. laevis adsl.L* RNA* on *myod1* expression at stage 12.5 p. 13
**Figure S13.** Expression of the myogenic regulatory factor *myod1* gene in the paraxial mesoderm is strongly affected by knock-down of *ppat* and *hprt1.L* genes p. 14
**Figure S14.** Statistical analysis of the effects associated with *adsl.L* knock-down on *myogenin* expression in craniofacial muscles in late tailbud stage embryos p. 15
**Figure S15.** Statistical analysis of the effects consecutive to *adsl.L* knock-down on *myod1* expression in hypaxial muscles in late tailbud stage embryos p. 15

### Supplementary Tables

**Table S1:** Yeast strains p. 16
**Table S2:** Plasmids used for functional complementation in yeast p. 17
**Table S3**: Plasmids used for mRNA synthesis p. 17
**Table S4**: Ribonucleotides probes used for *in situ* hybridization p. 18
**Table S5:** Oligonucleotides used for RT-PCR analyses p. 19
**Table S6:** Comparison of the purine biosynthesis pathways encoding genes and proteins between, *X. laevis, H. sapiens* and *X. tropicalis* p. 20-22
**Table S7:** Comparison of the purine biosynthesis pathways encoding genes and proteins between *X. laevis* and *S. cerevisiae* p. 23-25

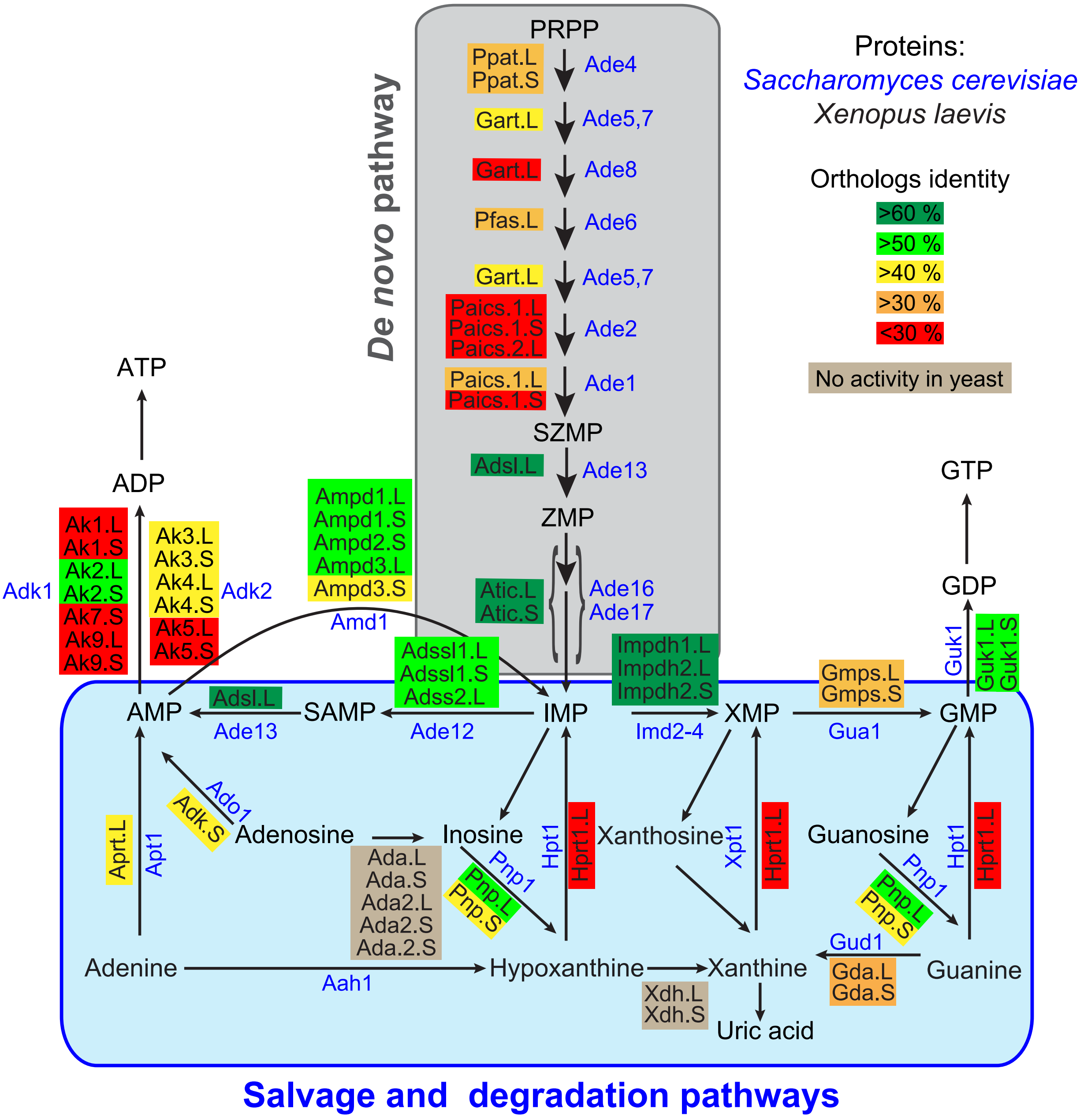

**Figure S1**. Conservation of purine biosynthesis pathways between the yeast *Saccharomyces cerevisia*e and *Xenopus laevis*. Features of the *X. laevis* and yeast enzymes are listed in Table S7. Abbreviations: AMP: adenosine monophosphate; GMP: guanosine monophosphate; IMP: Inosine monophosphate; PRPP: Phosphoribosyl pyrophosphate; SAMP: Succinyl-AMP; SZMP: Succinyl Amino Imidazole Carboxamide Ribonucleotide monophosphate; XMP: Xanthosine monophosphate; ZMP: Amino Imidazole CarboxAmide Ribonucleotide monophosphate.

```
ADSL    Hs     1-MAAGGDHGSPDS--------------------YRSPLASRYASPEMCFVFSDRYKFRTWRQLWLWLAEAEQTLGLPITDEQIQEMKSNLENI-72
               1-M..G....S.DS--------------------YRSPL.SRYAS.EM.F.FSD..KF.TWR.LWLWLA.AE..LGLPIT.EQIQEM..NLENI
Adsl.L  Xl     1-ME-GSSGLSMDSNTRITGSVSPLTPGPEEVMRYRSPLVSRYASREMAFNFSDSKKFQTWRRLWLWLAQAERSLGLPITEEQIQEMEANLENI-91

ADSL    Hs    73-DFKMAAEEEKRLRHDVMAHVHTFGHCCPKAAGIIHLGATSCYVGDNTDLIILRNALDLLLPKLARVISRLADFAKERASLPTLGFTHFQPAQ-164
                 DFKMAAEEEKRLRHDVMAHVHTF.HCCPKAA..IHLGATSCYVGDNTDLI.LR...DLLLPKLARV..RLADFA...A..PTLGFTH.QPAQ
Adsl.L  Xl    92-DFKMAAEEEKRLRHDVMAHVHTFAHCCPKAAPVIHLGATSCYVGDNTDLIVLRDGFDLLLPKLARVLNRLADFAEKYAEMPTLGFTHYQPAQ-183

ADSL    Hs   165-LTTVGKRCCLWIQDLCMDLQNLKRVRDDLRFRGVKGTTGTQASFLQLFEGDDHKVEQLDKMVTEKAGFKRAFIITGQTYTRKVDIEVLSVLA-256
                 LTTVGKR.CLW.QDLCMDL..L.R.R..LRFRGVKGTTGTQASFLQLF.GD..KVE.LD.MVT..AGFKRA.I.TGQTY.RKVD.EV.SVLA
Adsl.L  Xl   184-LTTVGKRACLWLQDLCMDLRNLERARNELRFRGVKGTTGTQASFLQLFDGDHDKVEELDRMVTSMAGFKRAYIVTGQTYSRKVDVEVVSVLA-275

ADSL    Hs   257-SLGASVHKICTDIRLLANLKEMEEPFEKQQIGSSAMPYKRNPMRSERCCSLARHLMTLVMDPLQTASVQWFERTLDDSANRRICLAEAFLTA-348
                 SLGA.VHKICTDIRLLANLKE.EEPFEK.QIGSSAMPYKRNPMRSERCCSLARHLMTL.M.PLQTASVQWFERTLDDSANRRICLAEAFLTA
Adsl.L  Xl   276-SLGATVHKICTDIRLLANLKELEEPFEKEQIGSSAMPYKRNPMRSERCCSLARHLMTLIMNPLQTASVQWFERTLDDSANRRICLAEAFLTA-367

ADSL    Hs   349-DTILNTLQNISEGLVVYPKVIERRIRQELPFMATENIIMAMVKAGGSRQDCHEKIRVLSQQAASVVKQEGGDNDLIERIQVDAYFSPIHSQL-440
                 D ILL.TLQNISEGLVVYPKVIERRIRQELPFMATENIIMAMVK.GG.RQDCHE.IRVLSQQA..VVKQEGGDNDLI.RIQ.D.YF.P.HA.H
Adsl.L  Xl   368-DIILSTLQNISEGLVVYPKVIERRIRQELPFMATENIIMAMVKNGGNRQDCHERIRVLSQQAGAVVKQEGGDNDLIFRIQSDSYFAPIHAHL-459

ADSL    Hs   441-FSPIHSQLDHLLDPSSFTGRASQQVQRFLEEEVYPLLKPYESVMKVKAELCL-484
                 F.PIH..L..LLDP.SF.GRA.QQV..FL.EEV.PLL.PY.S.M.VK.EL.L
Adsl.L  Xl   460-FAPIHAHLEQLLDPKSFIGRAPQQVLKFLKEEVIPLLSPYQSKMDVKMELEL-503
```

**Figure S2**. Sequence comparison between the *H. sapiens* (*Hs*) and *X. laevis* (*Xl*) adenylosuccinate lyase enzymes. Conserved amino acids are indicated in bold. Red and orange boxes point to residues required for catalysis and yellow and orange boxes to mutated residues found in *ADSL*-deficient patients.

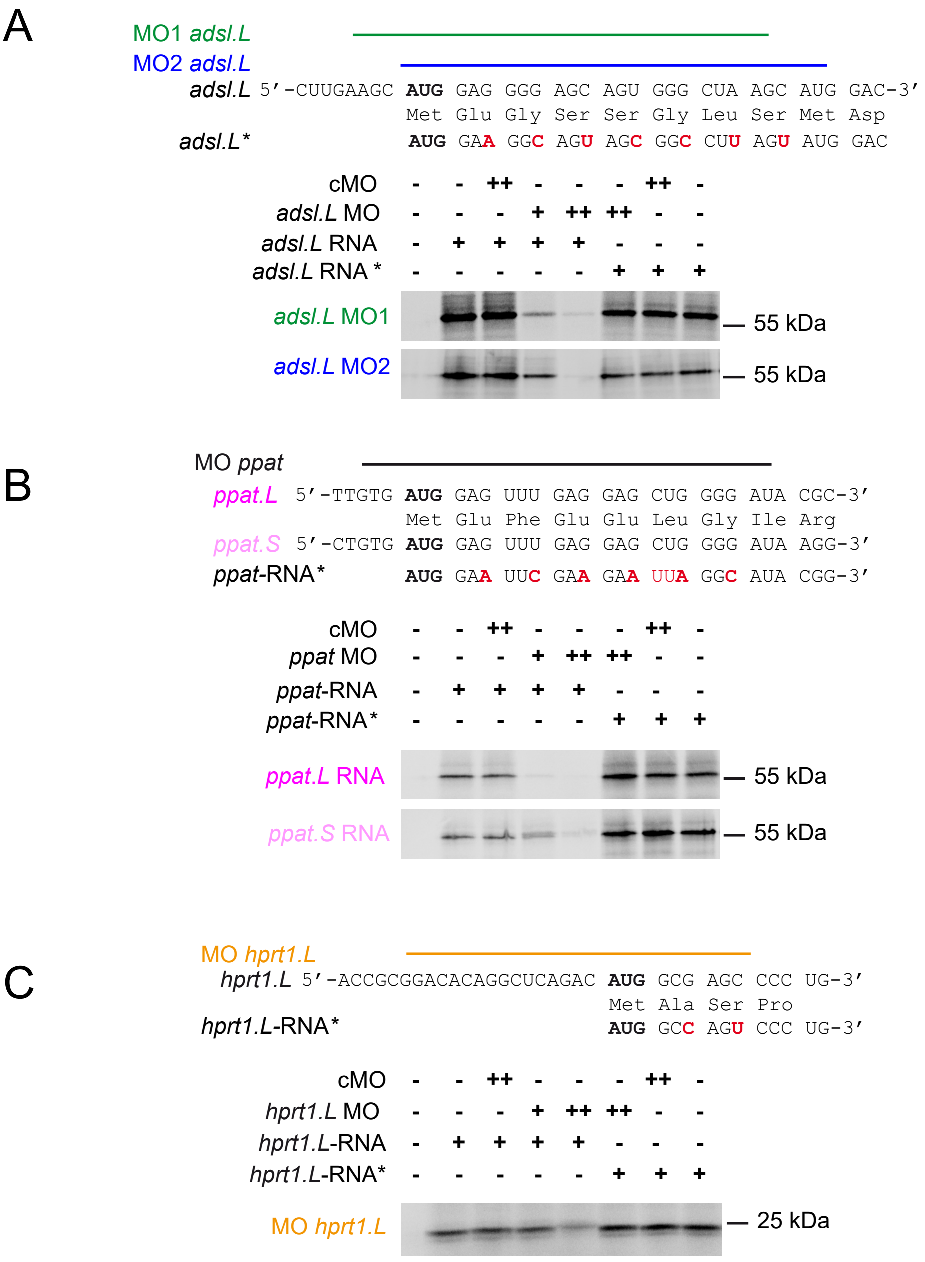

**Figure S3.** Validation of the morpholinos translation interference by *in vitro* translation. The *adsl.L* (**A**), *ppat.L/ppat.S* (**B**) and *hprt1.L* (**C**) RNAs were produced using the *mMessage mMachine SP6* kit (ambion) according to the supplier procedure. *In vitro* translation was then performed by using the reticulocyte rabbit lysate system kit (Promega) in 25 µl reaction mix containing 500 ng of indicated RNA, 0.5 µl of the amino acid mix w/o Met (1 mM), 2 µl of [$^{35}$S]-methionine (1,200 Ci/mmol; 10 mCi/ml), 17.5 µl of the reticulocytes lysate, 0.5 µL of RNAse inhibitor (40 u/µl; Promega) and in the presence or the absence of 40 (+) or 400 (++) ng of indicated Morpholino (MO). Proteins were separated by SDS-PAGE and radiolabeled proteins were detected by phosphorimaging (Typhoon biomolecular imager, Amersham). RNA* refers to mutated RNAs whose translation is not affected by the gene-specific MOs.

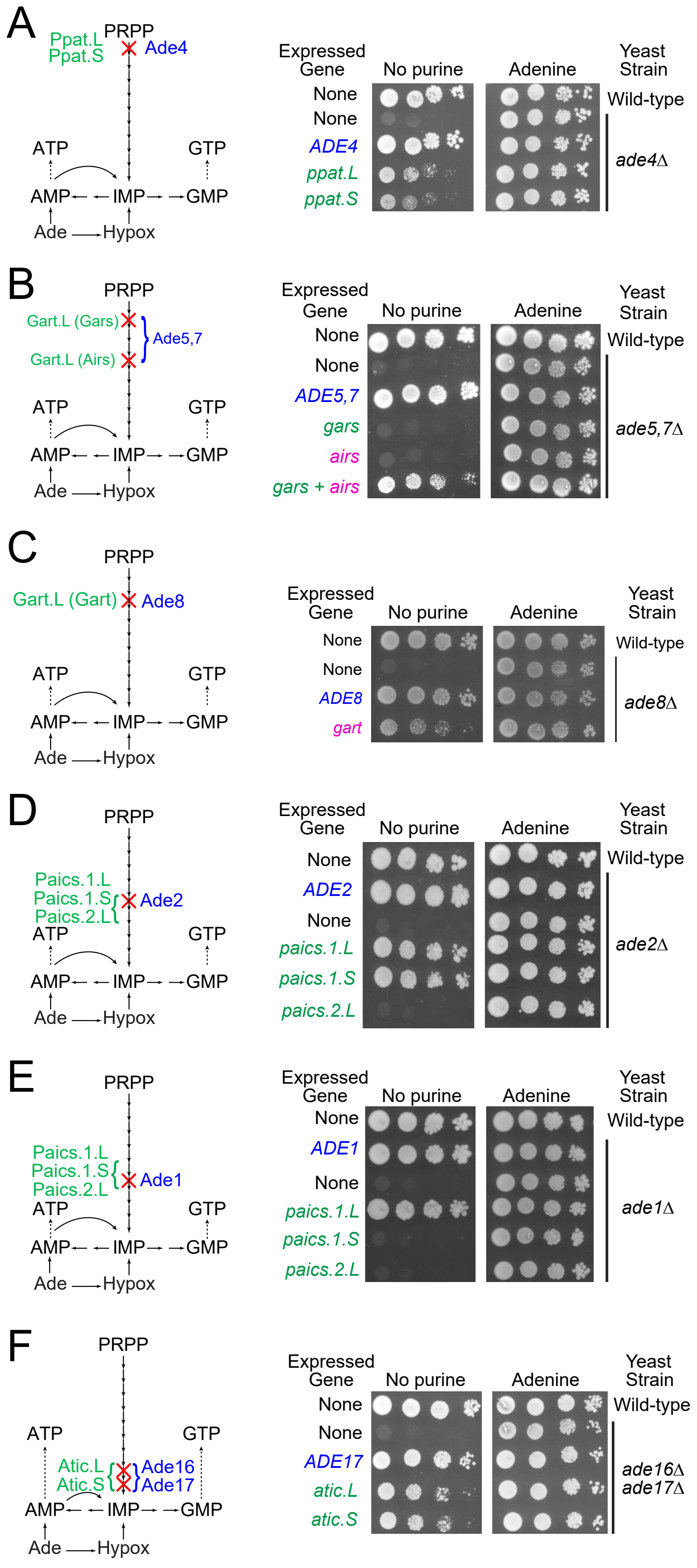

**Figure S4.** Functional complementation of yeast purine *de novo* pathway mutants by the *X. laevis* ortholog genes. Red crosses indicate the purine pathway steps knocked-out in each yeast mutant. *S. cerevisiae*, *X. laevis* and *X. tropicalis* protein or gene names are written in blue, green and pink, respectively. For Airs and Gart activities, functional complementation was performed by expressing the *X. tropicalis* corresponding open reading frames. Abbreviations: Ade: adenine; Airs; aminoimidazole ribonucleotide synthetase activity; Gars: glycinamide ribonucleotide synthetase activity; Gart: glycinamide ribonucleotide transformylase activity; Hypox: hypoxanthine; IMP: Inosine monophosphate; PRPP: Phosphorybosyl pyrophosphate. Yeast mutants were transformed with plasmids allowing expression of the indicated yeast (blue) or *X. laevis* (green) or *X. tropicalis* (pink) genes or with the empty vector (None). Transformants were serial (1/10) diluted and spotted on SDcasaW medium supplemented or not (no purine) with adenine as sole external purine source. Plates were imaged after 2 (**F**), 4 (**A**, **D-E**) or 7 days (**B-C**) at either 30°C (**A**, **D-F**) or 37°C (**B-C**).

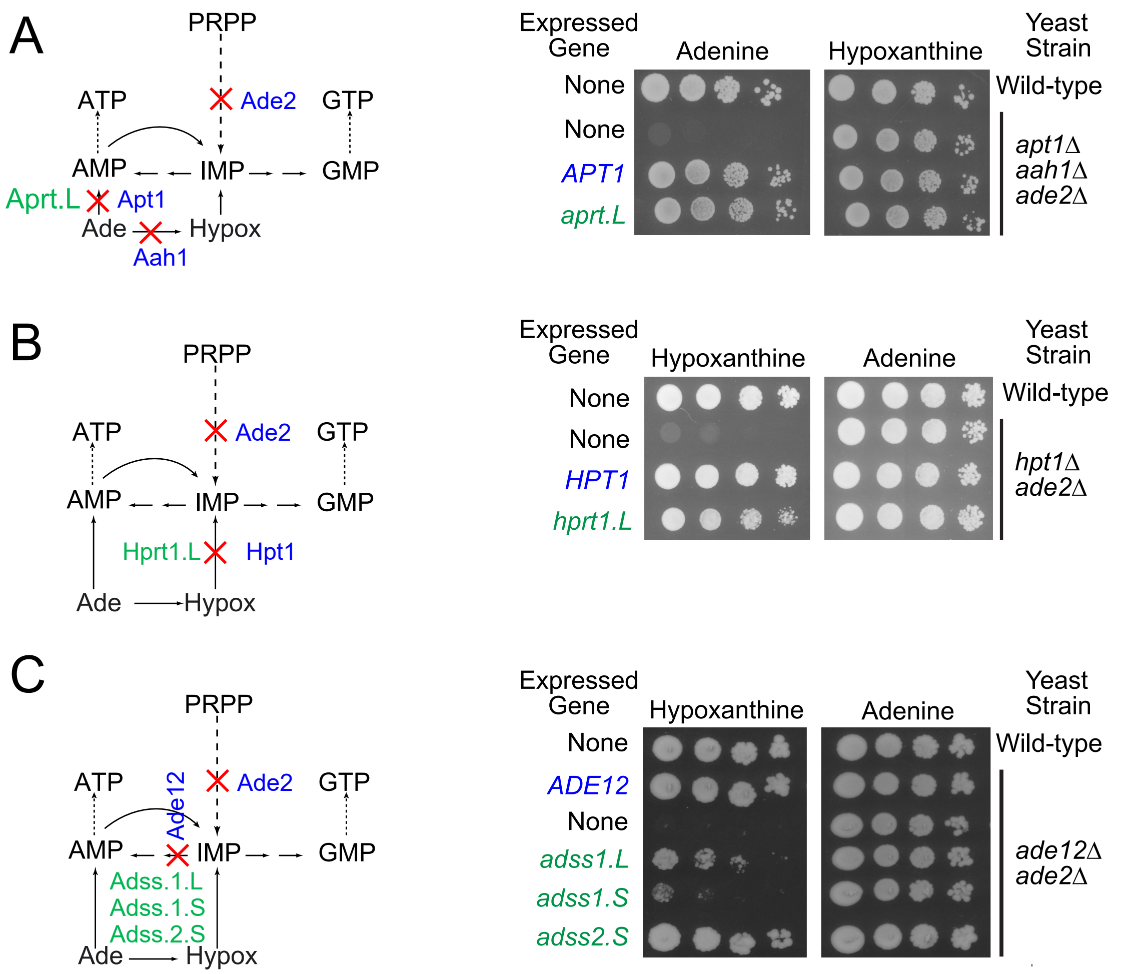

**Figure S5.** Functional complementation of yeast purine salvage knocked-out mutants by the *X. laevis* ortholog genes. The vertical dashed line symbolizes the *de novo* purine pathway. Red crosses are used to indicate the purine pathway steps absent in each yeast knock-out mutant. *S. cerevisiae* and *X. laevis* protein or gene names are written in blue and green, respectively. Ade: Adenine; Hypox: Hypoxanthine; IMP: Inosine monophosphate; PRPP: 5-phosphorybosyl-pyrophosphate. Wild-type and mutant yeast strains were transformed with plasmids allowing expression of the indicated *S. cerevisiae* (blue) or *X. laevis* (green) genes or with the empty vector (None). Transformants were serial diluted (1/10) and spotted on SDcasaW medium supplemented with either adenine or hypoxanthine as sole external purine source. Plates were imaged after 2 days at 30°C.

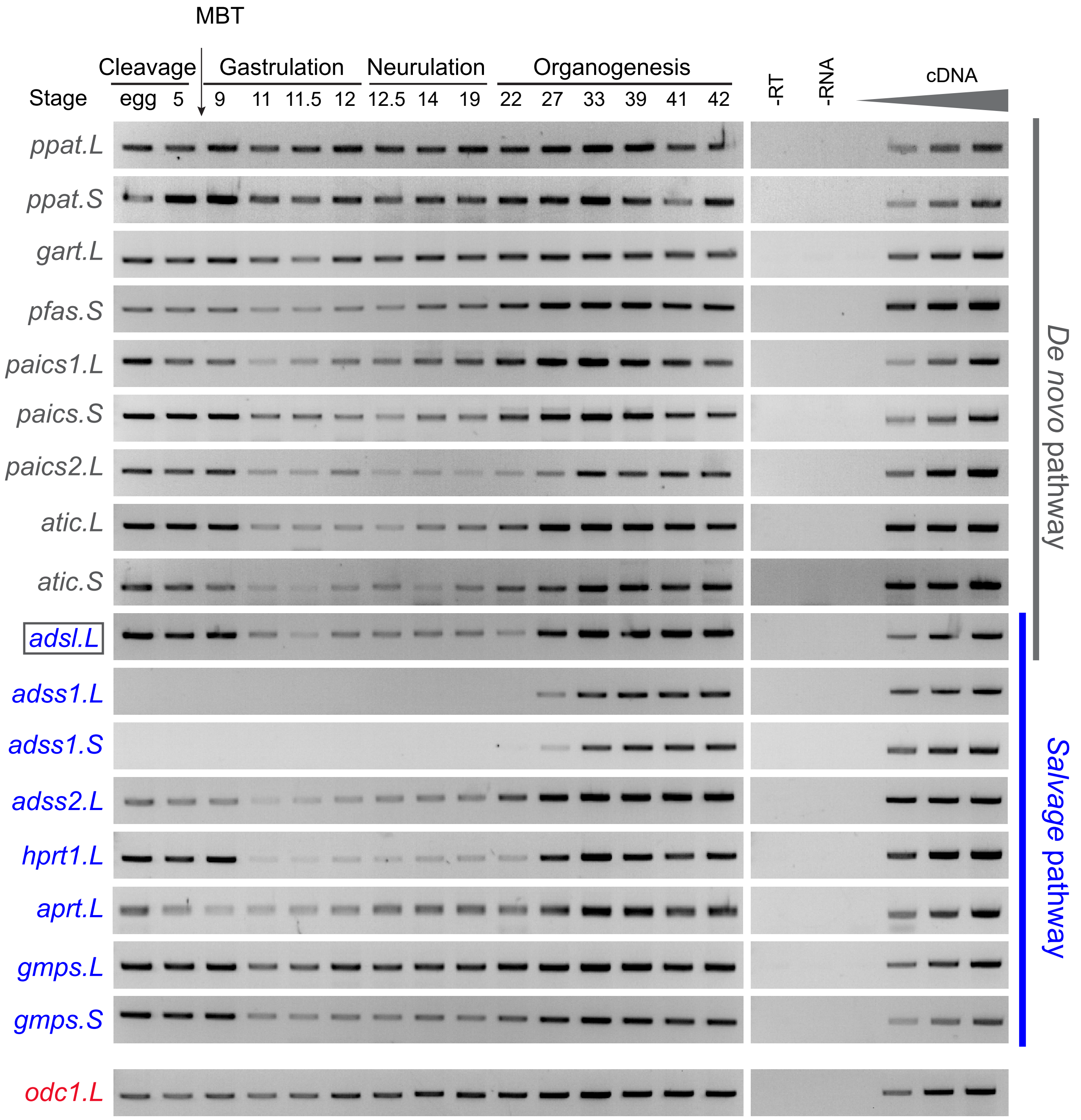

**Figure S6.** Temporal expression profiles of purine pathway genes during embryogenesis. The expression profile was determined for each gene by RT-PCR from cDNA of fertilized egg and whole embryo at different stages covering the different phases of *X. laevis* embryogenesis. Negative controls were performed in the absence of either reverse transcriptase (-RT) or total RNA extract (-RNA). The linearity was performed with doubling dilutions of cDNA. A loading control was done using *odc1.L* gene (in red). Names corresponding to genes encoding enzymes of the purine *de novo* and salvage pathways are written in grey and blue, respectively. The *adsl.L* gene encoding the adenylosuccinate lyase involved in both purine pathways is boxed in grey. Experiments were done at least twice ($n \geq 2$) on embryos obtained from *in vitro* fertilization of oocytes from two different females (N=2). Sense and antisense primers are described in Table S5. MBT: mid blastula transition.

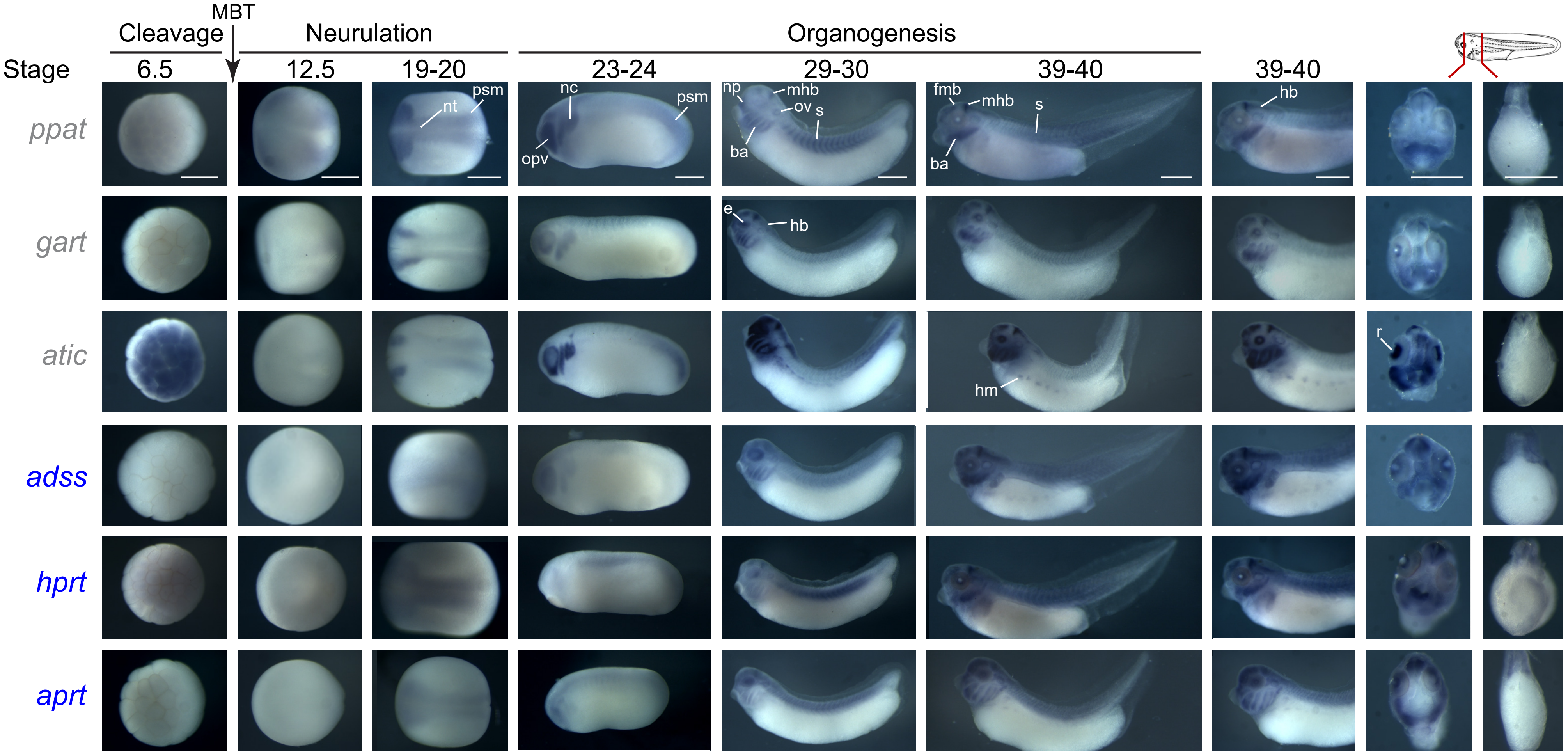

**Figure S7.** The expression profile was determined for each gene by *in situ* hybridization using antisense riboprobes (Table S4). No unspecific staining was detected with the control sense probes (Figure S8). MBT: Mid blastula transition. Grey and blue gene name colors refer to *de novo* and salvage purine pathways, respectively. Representative embryos were photographed. Stage 6.5: animal pole view, later stages: lateral views, with dorsal is up and anterior is left. Transverse section: dorsal is up. Abbreviations: ba, branchial arches; e, eye; fmb, forebrain-midbrain boundary; hb, hindbrain; hm, hypaxial muscles; l, lens; mhb, midbrain-hindbrain boundary; n: nasal placode; nc, neural crest; np, neural plate; nt, neural tube; opv, optical vesicle; ov, otic vesicle; psm, presomitic mesoderm; r, retina; s, somites. Bars: 0.5 mm.

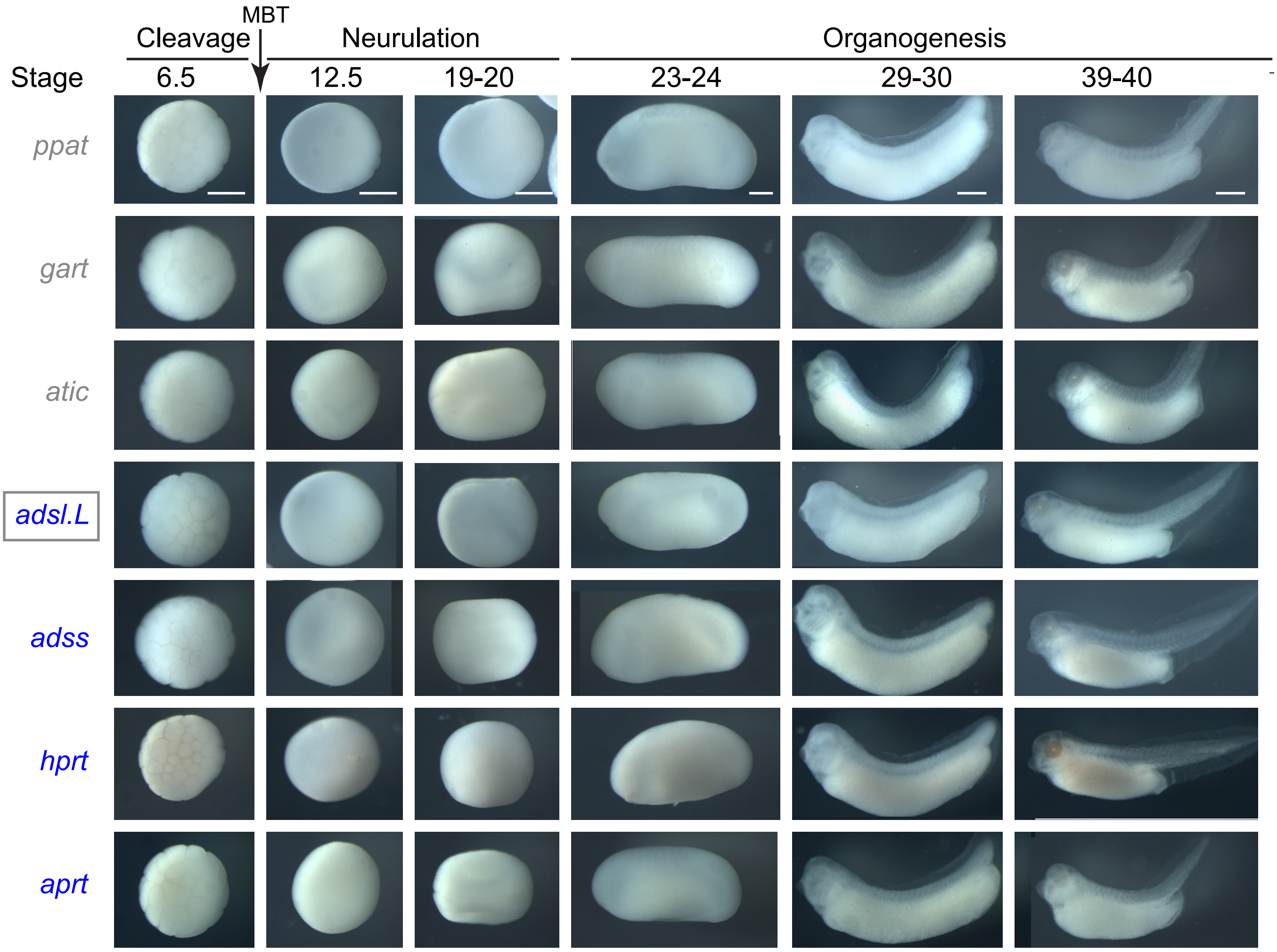

**Figure S8.** *In situ* hybridization using control sense riboprobes. *In situ* hybridization sense probes are described in Table S4. MBT: Mid blastula transition. Grey and blue gene name colors refer to *de novo* and salvage purine pathways, respectively. Representative embryos were photographed. Stage 6.5: animal pole view, later stages: lateral views, with dorsal is up and anterior is left. Bars: 0.5 mm.

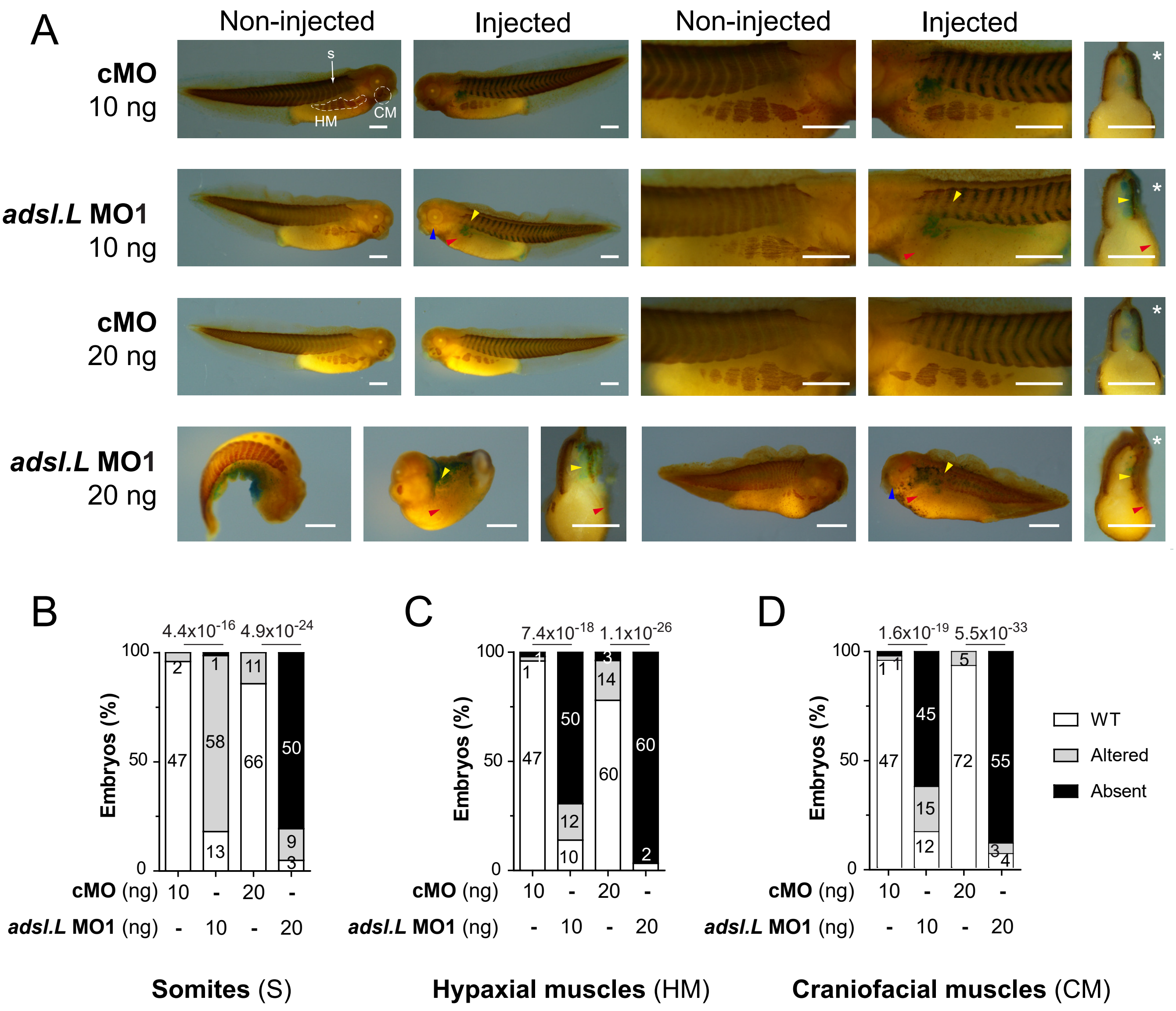

**Figure S9.** Severity of *adsl.L* knock-down-associated phenotypes is morpholino dose-dependent. 12-101 immunolabelling revealed a strong alteration of myogenesis and somitogenesis (**A-B**), hypaxial (**A**, **C**) and craniofacial (**A**, **D**) muscle formation in *adsl.L* morphant embryos. Representative images (**A**), quantification (embryo numbers in bars) and statistics of somite (**B**), hypaxial muscle (**C**) and craniofacial muscle (quadratoangularis + levator mandibularis longus) (**D**) phenotypes at tadpole stage. Yellow, blue and red arrowheads points to regions of somitic (S), craniofacial (CM) and hypaxial (HM) muscle defaults, respectively. Asterisks show injected side. Bars: 0.5 mm. Numbers above the bars in the histograms correspond to p-values.

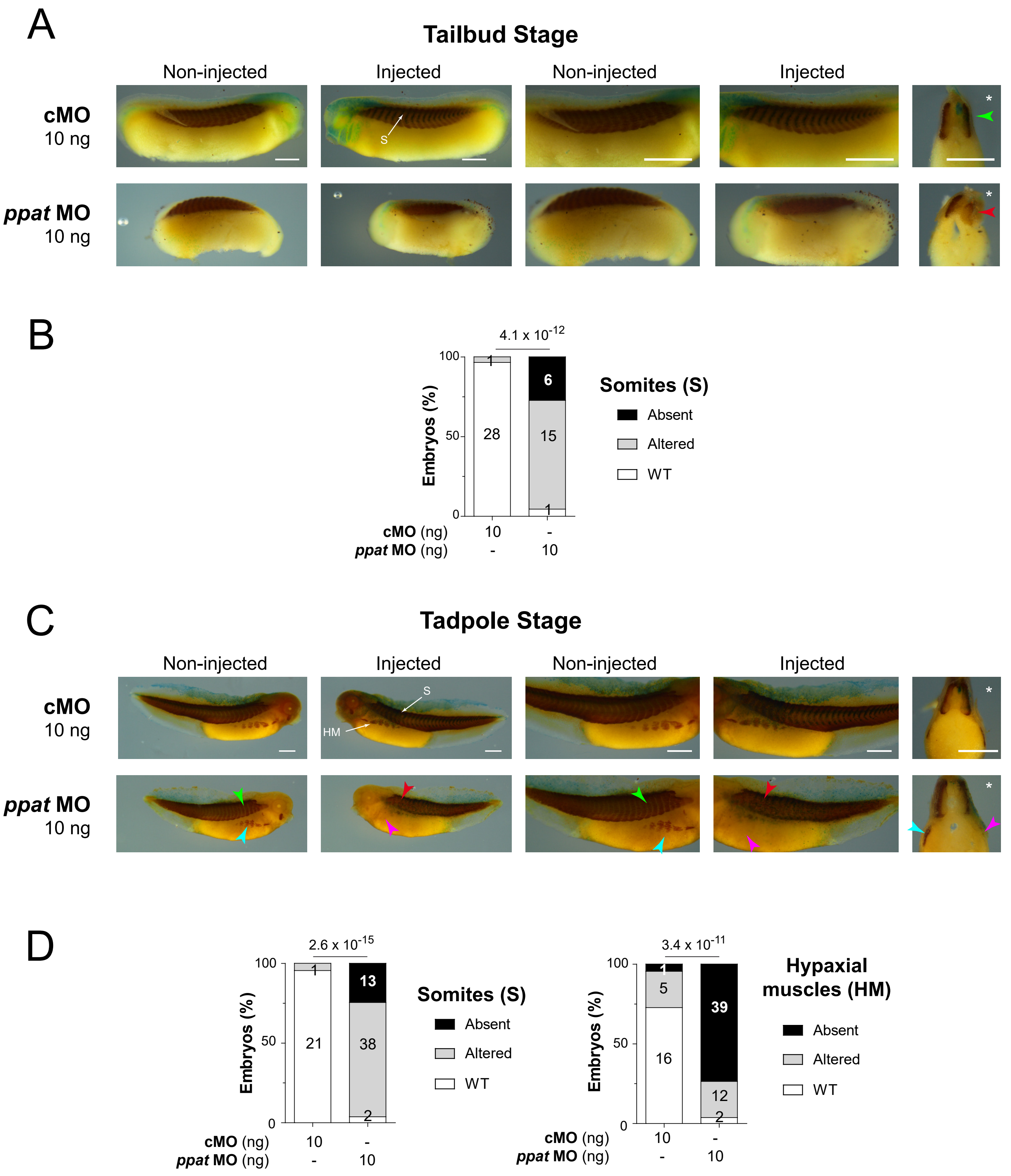

**Figure S10.** The *ppat.L* and *ppat.S* genes are required for somite, myotome and hypaxial muscle formation in *Xenopus laevis*. (**A-D**) A strong muscle alteration in *ppat* knock-down embryos is revealed by 12-101 immunolabelling. Representative images (**A, C**), quantification (embryo numbers in bars) and statistics (**B, D**) of somite and hypaxial muscle phenotypes at tailbud (**A-B**) and tadpole (**C-D**) stages. Green and red arrowheads point to normal v-shaped and altered somites, respectively; blue and pink arrowheads show normal and reduced 12/101 positive hypaxial muscle area, respectively. Injected side is indicated by asterisks. S: somites; HM: Hypaxial muscles. Bars: 0.5 mm. Numbers above the bars in the histograms correspond to p-values.

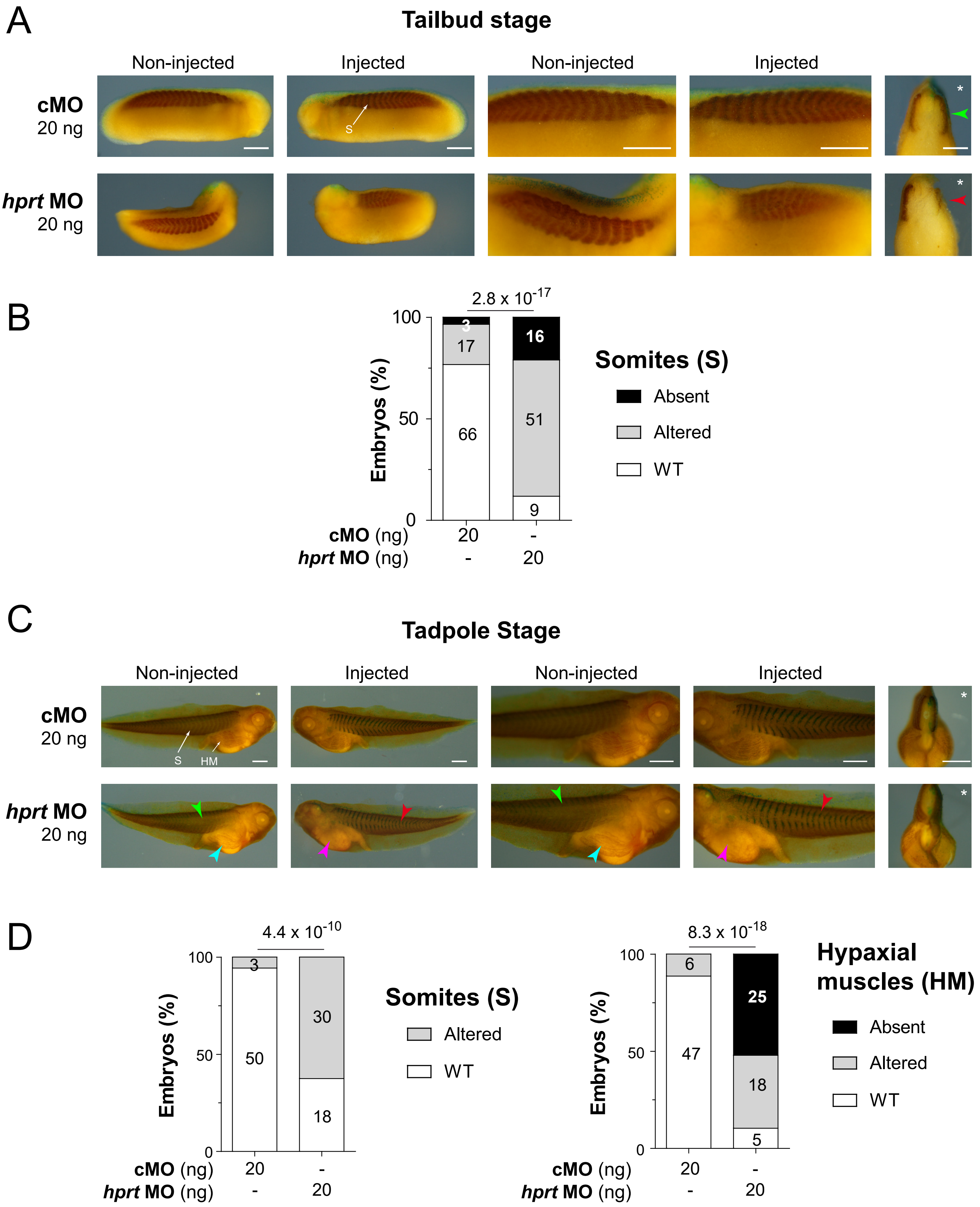

**Figure S11.** The *hprt1.L* gene is required for somite, myotome and hypaxial muscle formation in *Xenopus laevis*. (**A-D**) A strong muscle alteration is revealed by 12-101 immunolabelling in *hprt1.L* knock-down embryos. Representative images (**A, C**), quantification (embryo numbers in bars) and statistics (**B**, **D**) of somite and hypaxial muscle phenotypes at tailbud (**A-B**) and tadpole (**C-D**) stages. Green and red arrowheads point to normal v-shaped and altered somites, respectively; blue and pink arrowheads show normal and reduced 12/101 positive hypaxial muscle area, respectively. Injected side is indicated by asterisks. S: somites; HM: Hypaxial muscles. Bars: 0.5 mm. Numbers above the bars in the histograms correspond to p-values.

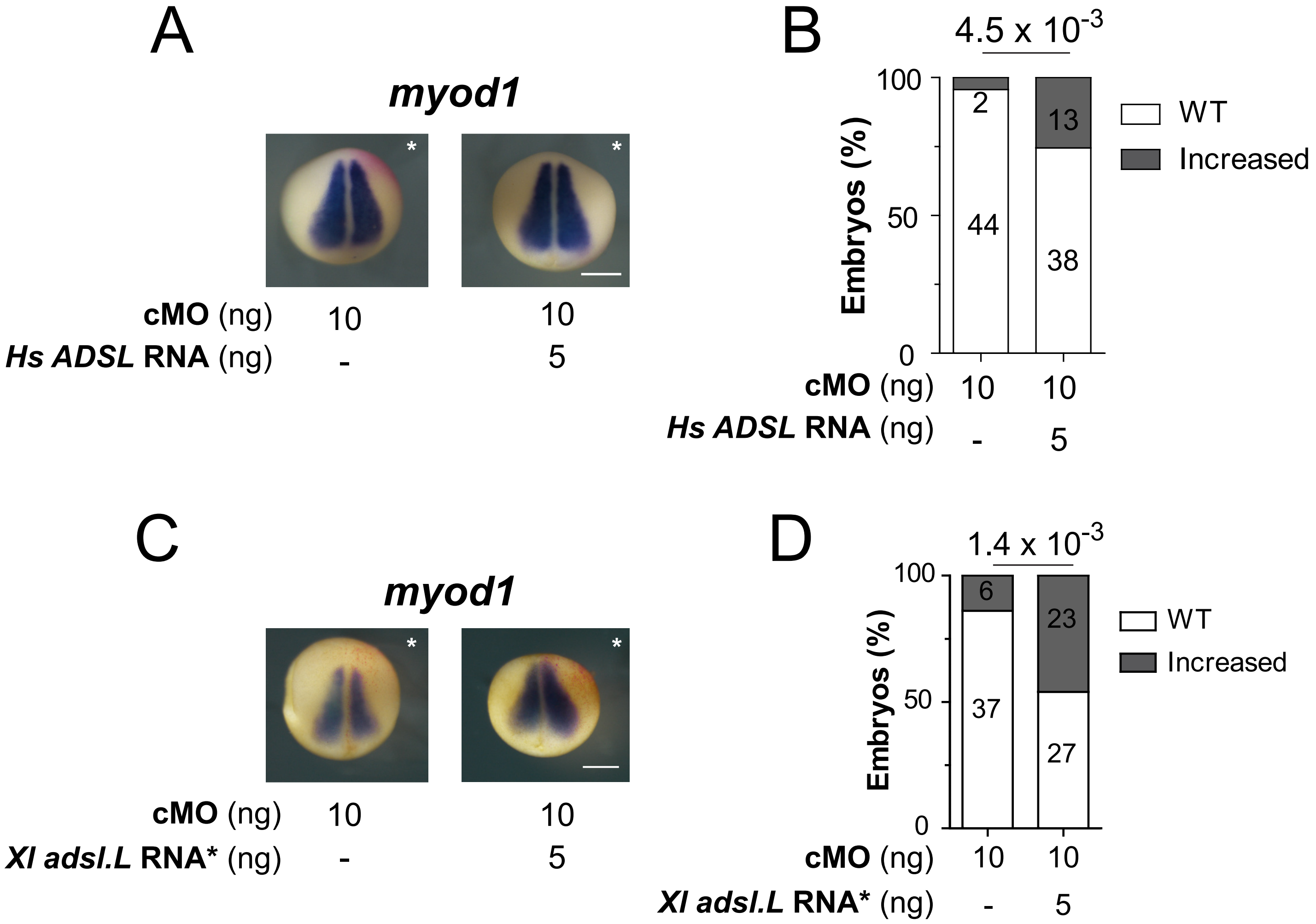

**Figure S12.** Effect of *H. sapiens ADSL* and *X. laevis adsl.L* RNA* on *myod1* expression at stage 12.5. (**A, C**) Expression of *myod1* was monitored by *in situ* hybridization at stage 12.5 on embryos injected with the control morpholino (cMO) and co-injected or not (-) with either the *H. sapiens ADSL* RNA (**A**) or the *X. laevis* MO non-targeted *adsl.L* RNA* (**C**), (**B, D**) Quantification (numbers in bars) and statistics of the *myod1* expression phenotypes are presented in (**A, C**). Injected side is indicated by asterisks. Bars: 0.5 mm. Numbers above the bars in the histograms correspond to p-values.

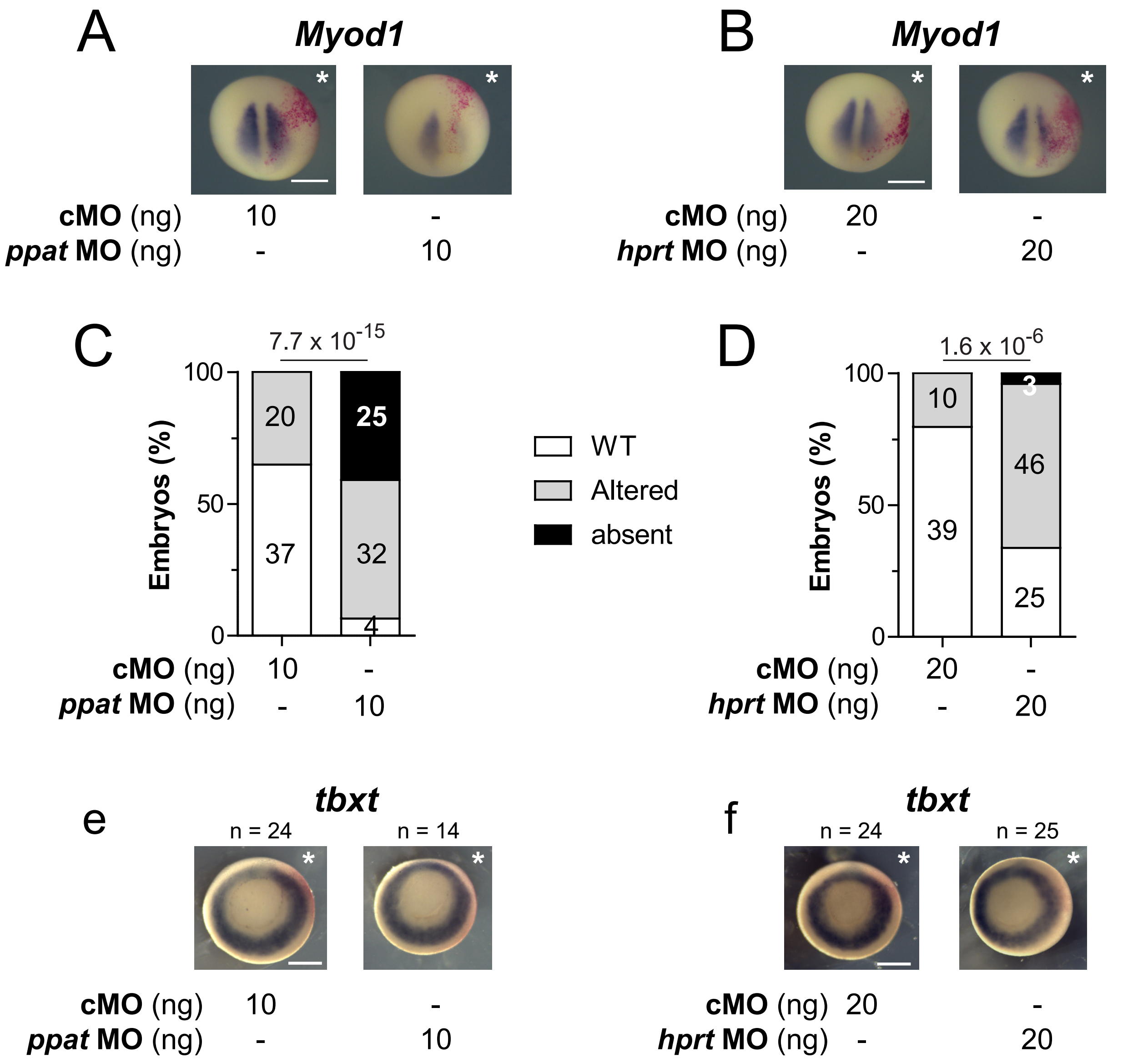

**Figure S13.** Expression of the myogenic regulatory factor *myod1* gene in the paraxial mesoderm is strongly affected by knock-down of *ppat* and *hprt1.L* genes. (**A-B**) Representative images of *myod1* expression alteration by the knock-down of either *ppat.L*/*ppat.S* (**A**) or *hprt1.L* (**B**) genes at stage 12.5, (**C-D**) Quantification (embryo numbers in bars) and statistics of the *myod1* expression phenotypes presented in (**A-B**). (**E-F**) Knock-down of both *ppat.L*/*ppat.S* (**E**) or *hprt1.L* (**F**) genes has no significant effect on general mesoderm formation, as revealed by the pan-mesoderm *tbxt* (*xbra*) expression pattern at stage 11. Injected side is indicated by asterisks. Bars: 0.5 mm. Numbers above the bars in the histograms correspond to p-values.

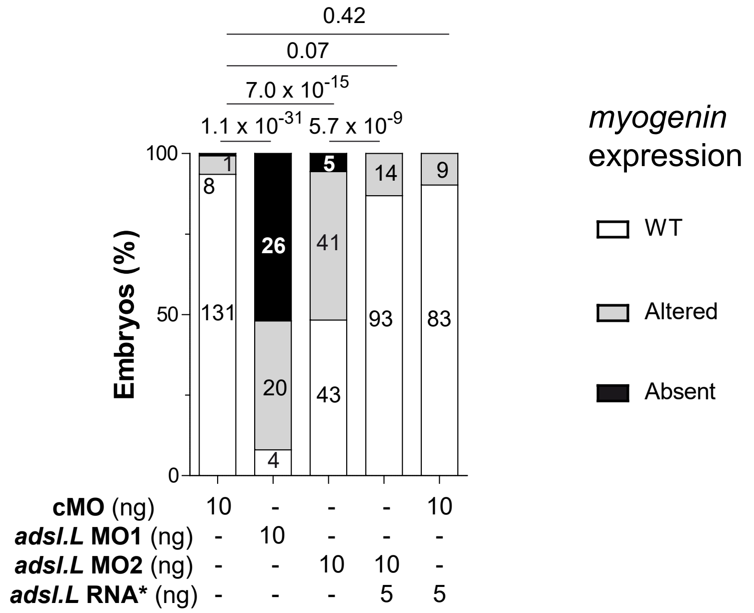

**Craniofacial muscles** (CM)

**Figure S14.** Statistical analysis of the effects associated with *adsl.L* knock-down on *myogenin* expression in craniofacial muscles in late tailbud stage embryos. Quantification (numbers in bars) and statistics of the *myogenin* expression phenotypes presented in Figure 6A. Numbers above the bars in the histograms correspond to p-values.

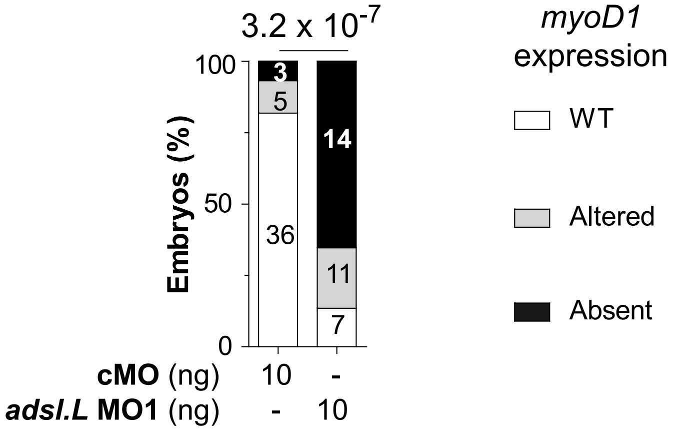

**Hypaxial muscles**
(HM)

**Figure S15.** Statistical analysis of the effects consecutive to *adsl.L* knock-down on *myod1* expression in hypaxial muscles in late tailbud stage embryos. Quantification (numbers in bars) and statistics of the *myod1* expression phenotypes presented in Figure 5C. Numbers above the bars in the histograms correspond to p-values.

**Table S1:** Yeast strains

| Strain | Genotype | Reference |
|---|---|---|
| BY4741 | *Matα ura3Δ leu2Δ his3Δ lys2Δ* | Euroscarf |
| BY4742 | *Mat***a** *ura3Δ leu2Δ his3Δ met15Δ* | Euroscarf |
| DS1-2B/1 | *Matα ura3Δ ade2 apt1 aah1* | R. Woods |
| Y1036 | *Matα ura3Δ leu2Δ his3Δ lys2Δ ade1::kanMX4* | Lab collection |
| Y1057 | *Matα ura3Δ leu2Δ his3Δ lys2Δ ade4::kanMX4* | Lab collection |
| Y1059 | *Matα ura3Δ leu2Δ his3Δ lys2Δ ade5,7::kanMX4* | Lab collection |
| Y1063 | *Matα ura3Δ leu2Δ his3Δ lys2Δ ade8::kanMX4* | Lab collection |
| Y1095 | *Matα ura3Δ leu2Δ his3Δ lys2Δ ade16::kanMX4 ade17::kanM*[illegible] | Lab collection |
| Y1133 | *Mat*a *ura3Δ leu2Δ his3Δ lys2Δ ade8::kanMX4* | Lab collection |
| Y3574 | *Matα ura3Δ leu2Δ his3Δ ade1::kanMX4 ade13::kanMX4* | Lab collection |
| Y8093 | *Matα ura3Δ leu2Δ his Δ ade2::kanMX4 hpt1::kanMX4* | Lab collection |
| Y11114 | *Matα ura3Δ leu2Δ his3Δ lys2Δ ade2::KanMX4 ade12::HIS3* | Lab collection |

**Table S2:** Plasmids used for functional complementation in *S. cerevisiae*

| Plasmid | I.M.A.G.E Clone used for amplification | Characteristics | Reference |
|---|---|---|---|
| pCM189 | N/A | *CEN ARS URA3 tet-OFF promoter* | Gari 1997[a] |
| p4930 | IRBHp990G1167D | *CEN ARS URA3 tet-atic.L. (X. laevis)* | This study |
| p4933 | IRBHp990H0610D | *CEN ARS URA3 tet-adsl.L. (X. laevis)* | This study |
| p5153 | IRAKp961G14156Q | *CEN ARS URA3 tet-atic.S. (X. laevis)* | This study |
| p5255 | IRBHp990F0459D | *CEN ARS URA3 tet-paics.1.L. (X. laevis)* | This study |
| p5257 | IRBHp990G071D | *CEN ARS URA3 tet-paics.1.S. (X. laevis)* | This study |
| p5303 | IRBHp990C0135D | *CEN ARS URA3 tet-adss.1.S. (X. laevis)* | This study |
| p5304 | IRBHp990B1280D | *CEN ARS URA3 tet-adss.1.L. (X. laevis)* | This study |
| P5318 | IRBH990B02030D | *CEN ARS LEU2 tet-hprt.L (X. laevis)* | This study |
| p5321 | IRAKp961E17253Q | *CEN ARS URA3 tet-aprt.L (X. laevis)* | This study |
| p5551 | IRBHp990H1017D | *CEN ARS URA3 tet-ppat.L.(X. laevis)* | This study |
| p5400 | IMAGp998O09145830Q | *CEN ARS URA3 tet-ppat.S (X. laevis)* | This study |
| p5550 | IRBH990B02030D | *CEN ARS LEU2 tet-hprt.S (X. laevis)* | This study |
| p5697 | IRAKp961P06157Q | *CEN ARS URA3 tet-paics.2 (X. laevis)* | This study |
| p5740 | IMAGp998B1011965Q | *CEN ARS URA3 tet-gart (X. tropicalis)* | This study |
| P5744 | IMAGp998B1011965Q | *CEN ARS URA3 tet-gars (X. tropicalis)* | This study |
| P5269 | IRAKp961C16299Q | *CEN ARS LEU2 tet-airs (X. laevis)* | This study |

[a] E Garí, L Piedrafita, M Aldea, E Herrero A set of vectors with a tetracycline-regulatable promoter system for modulated gene expression in Saccharomyces cerevisiae **1997** *Yeast* 13(9):837-48. DOI: 10.1002/(SICI)1097-0061(199707)13:9<837::AID-YEA145>3.0.CO;2-T

**Table S3**: Plasmids used for capped mRNA synthesis.
Linear. Enzyme: restriction enzyme used for linearization. *Hs: Homo sapiens*. * refers to morpholinos non-targeted sequences.

| mRNA | Linear. Enzyme | Plasmids |
|---|---|---|
| *ads1.L* | *Xho*I | IRBHp990H0610D |
| *ads1.L** | *Xho*I | pBF-*adsl.L** from IRBHp990H0610D |
| *Hs ADSL* | *Not*I | pCS2$^{+}$-HsADSL |
| *ppat.L* | *Xba*I | IRBHp990H1017D |
| *ppat.L** | *Sal*I | pBF-*ppat.L** from IRBHp990H1017D |
| *ppat.S* | *Hind*III | IMAGp998O0914583Q |
| *ppat.S** | *Xho*I | pBF-*ppat.S** from IMAGp998O0914583Q |
| *hprt1.L* | *Not*I | IRBH990B02030D |
| *hprt1.L** | *Sac*I | pBF-*hprt1.L** from IRBH990B02030D |
| *LacZ* | *Xho*I | pSP6nucβgal |

**Table S4**: Ribonucleotides probes used for *in situ* hybridization.
CDS: coding sequence. Linear. Enzyme : restriction enzyme used for linearization. pSK: p-BlueScript plasmid. RNA Pol.: RNA polymerase. UTR: untranslated region

| Targeted genes | Probe | Linear. Enzyme | RNA Pol. | Plasmids |
|---|---|---|---|---|
| *adsl.L* | sense | *Kpn*I | T7 | p5265 *adsl.L*-pSK (CDS +3'UTR; 480 bp ) from IRBHp990H0610D |
| | antisense | *Sac*I | T3 | p5265 *adsl.L*-pSK (CDS +3'UTR; 480 bp ) from IRBHp990H0610D |
| *adss1.L* *adss1.S* | sense | *Sac*I | T3 | p5634 *adss1.L*-pSK (5'UTR + CDS; 444 bp ) from IRBHp990C0135D |
| | antisense | *Sal*I | T7 | p5634 *adss1.L*-pSK (5'UTR + CDS; 444 bp ) from IRBHp990C0135D |
| *adss2.L* | sense | *SacI* | T3 | p5428 *adss2.L*-pKS (5'UTR + CDS; 444 bp ) from IRBHp990E047D |
| | antisense | SalI | T7 | p5428 *adss2.L*-pKS (5'UTR + CDS; 444 bp ) from IRBHp990E047D |
| *aprt.L* | sense | *Kpn*I | T7 | p5435 *aprt.L*-pKS (CDS +3'UTR; 779 bp ) from IRAKp961E17253Q |
| | antisense | *Eco*RV | T3 | p5435 *aprt.L*-pKS (CDS +3'UTR; 779 bp ) from IRAKp961E17253Q |
| *atic.L* *atic.S* | sense | *Xho*I | SP6 | P4915 *atic.L*-pExpress1 (CDS +3'UTR; 1854 bp ) from IRBHp990G1167D |
| | antisense | *Eco*RI | T7 | P4915 *atic.L*- pExpress1 (CDS +3'UTR; 1854 bp ) from IRBHp990G1167D |
| *hprt.L* | sense | *Ase*I | T7 | p5454 *hprt.L*-pKS (5'UTR + CDS; 592 bp ) from IRBH990B02030D |
| | antisense | *Sac*I | T3 | p5454 *hprt.L*-pKS (5'UTR + CDS; 592 bp ) from IRBH990B02030D |
| *gart.L* | sense | *Bam*HI | T3 | p5426 *gart.L*-pKS (CDS +3'UTR ; 530 bp ) from IRAKp961C16299Q |
| | antisense | *Eco*RV | T7 | p5426 *gart.L*-pKS (CDS +3'UTR ; 530 bp ) from IRAKp961C16299Q |
| *ppat.L* *ppat.S* | sense | *Not*I | T3 | p5527 *ppat.L*-pSK (CDS ; 481 bp) from XL.29008 |
| | antisense | *Eco*RV | T7 | p5527 *ppat.L*-pSK (CDS ; 481 bp) from XL.29008 |

**Table S5:** Oligonucleotides used for RT-PCR analyses. Amplification temperature and number of RT-PCR cycles were determined to obtain a single PCR product.

| **Amplified gene** | **Sense oligonucleotide (5'-3')** | **Antisense oligonucleotide (5'-3')** | **Amplification Temperature (°C)** | **Nb of cycles** |
|---|---|---|---|---|
| *ppat.L* | GTTAGTCCCTGTGGCCGCT | GCCCATGCCCTTGTGCATTC | 54 | 28 |
| *ppat.S* | GAAGCGCGAGGTGTGTGTG | GCCCATGCCCTTGTGCATTC | 58 | 29 |
| *gart.L* | CAGAGACAGTTCTAGTGATTGG | GATAATCCCTGCTGCCAGAG | 56 | 29 |
| *pfas.S* | CTGACAGAACGTGCAGGG | CTAGTCTGGAACTTTATGAG | 53 | 29 |
| *paics.1.S* | CAGAACCACGTGGTACTGCC | GATCCCAGCCTCCTGCAGC | 56 | 30 |
| *paics.1.L* | GAAATGGAGTCTTACGCAGAAC | GATCCCAGCCTCCTGCAGC | 53 | 27 |
| *paics.2.L* | GCATTTGTTAAGAGGTGCAG | CCAAACTTAATCCTCATGTCC | 55 | 28 |
| *adsl.L* | ATGGCCTTCAACTTCAGCGA | AACGTTGGCATCTCTGCGTA | 55 | 28 |
| *atic.L* | GCTGCTAGCGACTTTATCCAG | GGGTACAGGTTACACACAAC | 57 | 31 |
| *atic.S* | GCTGTTTGCGGAGATGGAG | GGGTACAGGTTACACACAAC | 53 | 30 |
| *adss1.L* | CAAGTGGCATTATCAACCCC | CTTTGGATGAATATGTTGGTC | 51 | 30 |
| *adss1.S* | GTGGCATTATAAATCCTAAAGC | CTTTGGATGAATATGTTGGTC | 53 | 29 |
| *Adss2.L* | CCGCTACAGTAAGCGTAAC | GCTTTCATAGGAAATGGTGTG | 51 | 28 |
| *hprt1.L* | CTAAACATTATGCAGCCGATC | CATTCTTGCCTGTCAAGGTGG | 54 | 26 |
| *hprt1.S* | CTAAACACTACGCCGCCAGC | CATTCTTGCCTGTCAAGGTGG | 52 | 29 |
| *aprt.L* | CAGATTATGTCCGATCAGGAG | GCTAACAGATTCTGTGGGAC | 57 | 29 |
| *gmps.L* | GAGCGATGGGCAGAGATC | CTGTAAGAGCGACAGTCC | 56 | 28 |
| *gmps.S* | GAGGGTTGGGTAGAGAAC | CTGTAAGAGCGACAGTCC | 53 | 29 |
| *odc1.L* | GTCAATGATGGAGTGTATGGATC | TCCATTCCGCTCTCCTGACCAC | 55 | 23 |

**Table S6:** Comparison of the purine biosynthesis pathways encoding genes and proteins between, *X. laevis*, *H. sapiens* and *X. tropicalis*. Alignments were performed using https://blast.ncbi.nlm.nih.gov. * XB: xenbase: https://www.xenbase.org/entry/

| *Xenopus laevis* | | | *Homo sapiens* | | | | *Xenopus tropicalis* | | | | |
|---|---|---|---|---|---|---|---|---|---|---|---|
| **Protein name** | **Accession Number** | **XB* Gene ID** | **Protein name** | **Accession Number** | **Identity %** | **Coverage %** | **Protein name** | **Accession Number** | **Identity %** | **Coverage %** | **Enzymatic activity** |
| **Ada.L**<br>**Ada.S** | XP_018090401.1<br>NP_001085740.1 | 17336388<br>950506 | ADA | NP_000013.2 | 74<br>70 | 97<br>98 | Ada | NP_001011025.1 | 97 | 100 | Adenosine deaminase |
| **Ada2.L**<br>**Ada2.S** | NP_001090531.1<br>NP_001089165 | 6254244<br>6251688 | ADA2<br>ADA2 | NP_001269154.1 | 57<br>59 | 96<br>97 | Ada2<br>Ada2 | XP_031754454.1 | 90<br>88 | 99<br>100 | Adenosine deaminase |
| **Ada.2.S** | NP_001087740.1 | 5929096 | ADA.2 | No significant homolog | | | Ada.2 | NP_001107369.1 | 90 | 100 | Adenosine deaminase |
| **Adk.S** | NP_001086357.1 | 997231 | ADK | NP_001114.2 | 82 | 95 | Adk | NP_001016698 | 98 | 99 | Adenosine kinase |
| **Adsl.L** | NP_001080593 | 380285 | ADSL | NP_000017.1 | 83 | 93 | Adsl | NP_001005457.1 | 96 | 100 | Adenylosuccinate lyase |
| **Adssl1.L**<br>**Adssl1.S** | NP_001090012.1<br>NP_001087505.1 | 5758296<br>6254009 | ADSS1<br>ADSS1 | NP_689541.1 | 88<br>87 | 100<br>100 | Adssl1<br>Adssl1 | NP_001004939.1 | 97<br>96 | 100<br>100 | Adenylosuccinate synthase |
| **Adss2.L** | NP_001080088.1 | 944129 | ADSS2 | NP_001117.2 | 86 | 94 | Adss2 | NP_989047.1 | 95 | 100 | Adenylosuccinate synthase |
| **Ak1.L**<br>**Ak1.S** | NP_001087683.1<br>NP_001085451.1 | 6253770<br>6251725 | AK1<br>AK1 | NP_001305051.1 | 74<br>81 | 98<br>98 | Ak1<br>Ak1 | NP_001006817.1 | 91<br>100 | 97<br>100 | Adenylate kinase |
| **Ak2.L**<br>**Ak2.S** | XP_018102289.1<br>NP_001080232.1 | 17344313<br>6254479 | AK2<br>AK2 | NP_001616.1 | 82<br>79 | 100<br>100 | Ak2<br>Ak2 | XP_012812152.1 | 96<br>97 | 100<br>100 | Adenylate kinase |
| **Ak3.L**<br>**Ak3.S** | NP_001089446.1<br>NP_001084561.1 | 977031<br>17332462 | AK3<br>AK3 | NP_057366.2 | 74<br>75 | 97<br>97 | Ak3<br>Ak3 | XP_012812420.1 | 90<br>92 | 100<br>100 | Adenylate kinase |
| **Ak4.L**<br>**Ak4.S** | XP_018113718.1<br>XP_018116248.1 | 17341782<br>959957 | AK4 | NP_001005353.1 | 77<br>74 | 96<br>94 | Ak4<br>Ak4 | XP_002931643.1 | 96<br>97 | 100<br>100 | Adenylate kinase |
| **Ak5.L**<br>**Ak5.S** | XP_018113675.1<br>XP_018116212.1 | 6487811<br>17345311 | AK5<br>AK5 | AAH36666.1<br>NP_777283.1 | 72<br>71 | 100<br>100 | Ak5<br>Ak5 | XP_012815974.2 | 93<br>92 | 100<br>100 | Adenylate kinase |
| **Ak6.L**<br>**Ak6.S** | NP_001089528.1<br>NP_001087040.1 | 972297<br>17335043 | AK6<br>AK6 | NP_057367.1 | 76<br>78 | 100<br>100 | Ak6<br>Ak6 | NP_001017167.1 | 95<br>97 | 100<br>100 | Adenylate kinase |

| | | | | | | | | | | | |
|---|---|---|---|---|---|---|---|---|---|---|---|
| **Ak7.S** | NP_001081046.1 | 953552 | AK7 | NP_689540.2 | 63 | 99 | Ak7 | NP_001011352.1 | 93 | 100 | Adenylate kinase |
| **Ak8.L** | NP_001088862.1 | 5831351 | AK8 | NP_689785.1 | 55 | 98 | Ak8 | NP_989104.1 | 90 | 100 | Adenylate kinase |
| **Ak9.L** | XP_018118807 | 17335922 | AK9 | NP_001316531.1 | 52 | 34 | Ak9 | XP_031757751.1 | 85 | 96 | Adenylate kinase |
| **Ak9.S** | XP_018118808 | 17335923 | AK9 | | 52 | 34 | Ak9 | | 85 | 96 | |
| **Ampd1.L** | XP_018101841.1\| | 17335636 | AMPD1 | NP_000027.2 | 74 | 99 | Ampd1 | XP_002935821.2 | 95 | 100 | Adenosine monophosphate deaminase |
| **Ampd1.S** | XP_018104675.1 | 17335637 | AMPD1 | | 72 | 99 | Ampd1 | | 92 | 100 | |
| **Ampd2.S** | XP_018104829 | 6489067 | AMPD2 | NP_001355738.1 | 80 | 96 | Ampd2 | XP_031752724.1 | 96 | 100 | Adenosine monophosphate deaminase |
| **Ampd3.L** | XP_018112762.1 | 6489056 | AMPD3 | NP_000471.1 | 80 | 99 | Ampd3 | NP_001025687.1 | 94 | 100 | Adenosine monophosphate deaminase |
| **Ampd3.S** | XP_018115800.1 | 17344991 | AMPD3 | | 80 | 99 | Ampd3 | | 95 | 100 | |
| **Aprt.L** | XP_018096991.1 | 6253243 | APRT | NP_000476.1 | 65 | 100 | Aprt | NP_001007941.1 | 93 | 100 | Adenine phosphoribosyltransferase |
| **Atic.L** | NP_001090100.1 | 998812 | ATIC | NP_004035.2 | 78 | 99 | Atic | NP_001005460.1 | 95 | 100 | Amino-Imidazole CarboxAmide Ribonucleotide transformylase and IMP cyclohydrolase |
| **Atic.S** | XP_018094436.1 | 17331460 | ATIC | | 79 | 99 | Atic | | 96 | 100 | |
| **Gda.L** | NP_001083074.1 | 987047 | GAH | NP_004284.1 | 61 | 93 | Gda | XP_004910825.1 | 93 | 96 | Guanine deaminase |
| **Gda.S** | XP_018099342.1 | 17337222 | GAH | | 60 | 99 | Gda | | 83 | 99 | |
| **Gart.L** | NP_001093352.1 | 6251947 | GART | NP_000810.1 | 75 | 98 | Gart | XP_012813454.1 | 92 | 98 | Phosphoribosylglycinamide-formyltransferase, phosphoribosylglycinamide synthetase and phosphoribosylaminoimidazole synthetase |
| **Gmps.L** | XP_018119259.1 | 950699 | GMPS | NP_003866.1 | 92 | 98 | Gmps | XP_002933290.3 | 97 | 98 | Guanine monophosphate synthase |
| **Gmps.S** | XP_018121231.1 | 17340569 | GMPS | | 93 | 100 | Gmps | | 98 | 100 | |
| **Guk1.L** | NP_001087146.1 | 6254574 | GUK | NP_000849.1 | 70 | 97 | Guk1 | NP_001034818.1 | 89 | 100 | Guanylate kinase |
| **Guk1.S** | NP_001086807.1 | 17334557 | GUK | | 69 | 97 | Guk1 | | 90 | 100 | |
| **Hprt1.L** | NP_001090235.1 | 6078713 | HGPRT | NP_000185.1 | 89 | 99 | Hprt1 | NP_989312.1 | 100 | 100 | Hypoxanthine-Guanine phosphorybosyl transferase |
| **Impdh1.L** | NP_001080792 | 957455 | IMPDH1 | NP_000874.2 | 92 | 100 | Impdh1 | NP_001017283.1 | 98 | 100 | Inosine monophosphate dehydrogenase |
| **Impdh2.L** | NP_001083990.1 | 17345296 | IMPDH2 | NP_000875.2 | 93 | 100 | Impdh2 | NP_001008066.1 | 98 | 100 | Inosine monophosphate dehydrogenase |
| **Impdh2.S** | NP_001082410.1 | 478732 | IMPDH2 | | 93 | 100 | Impdh2 | | 97 | 100 | |
| **Paics.1.L** | NP_001080163 | 17334793 | PAICS | NP_001072992.1 | 81 | 100 | Paics.1 | XP_012811124.2 | 96 | 100 | Phosphoribosylaminoimidazole carboxylase Phosphoribosylaminoimidazolesuccino-carboxamide synthetase |
| **Paics.1.S** | NP_001086248.1 | 965998 | PAICS | | 82 | 98 | Paics.1 | | 95 | 100 | |

| | | | | | | | | | | | |
|---|---|---|---|---|---|---|---|---|---|---|---|
| **Paics.2.L** | XP_018085590.1 | 5872998 | PAICS | NP_001072992.1 | 57 | 37 | Paics.2 | NP_001090685.1 | 86 | 97 | Phosphoribosylaminoimidazole carboxylase<br>Phosphoribosylaminoimidazolesuccino-carboxamide synthetase |
| **Pfas.S** | XP_018094887.1 | 6485675 | PFAS | NP_036525.1 | 65 | 99 | Pfas | XP_012825646.2 | 93 | 99 | Phosphoribosylformylglycinamidine synthase |
| **Pnp.L** | NP_001079809.1 | 1011603 | PNP | NP_000261.2 | 68 | 95 | Pnp | NP_001006720.1 | 88 | 98 | Purine nucleoside phosphorylase |
| **Pnp.S** | XP_018099448.1 | 17337633 | PNP | | 66 | 93 | Pnp | | 90 | 100 | |
| **Ppat.L** | NP_001083491.1 | 972209 | GPAT | NP_002694.3 | 80 | 100 | Ppat | NP_989313.1 | 96 | 100 | Phosphoribosylpyrophosphate amidotransferase |
| **Ppat.S** | XP_041435416 | 17340761 | GPAT | | 79 | 99 | Ppat | | 96 | 100 | |
| **Xdh.L** | XP_018117710.1 | 6485774 | XDH | NP_000370.2 | 70 | 99 | Xdh | XP_031758260.1 | 89 | 100 | Xanthine dehydrogenase/oxidase |
| **Xdh.S** | XP_018120131.1 | 17331587 | XDH | | 70 | 92 | Xdh | | 90 | 100 | |

**Table S7:** Comparison of the purine biosynthesis pathways encoding genes and proteins between *X. laevis* and *S. cerevisiae*. Alignments were performed using https://blast.ncbi.nlm.nih.gov. * XB: xenbase: https://www.xenbase.org/entry/.
[a] No significant alignment between the yeast and *X. Laevis* Hprt1.L entire sequences.
ada: adenine deaminase; xdh: xanthine dehydrogenase/oxidase

| *Xenopus laevis* | | | *Saccharomyces cerevisiae* | | | | | |
|---|---|---|---|---|---|---|---|---|
| **Protein name** | **Accession Number** | **XB* Gene ID** | **Protein name** | **Accession Number** | **Identity %** | **SGD Locus ID** | **Coverage %** | **Enzymatic activity** |
| Ada.L<br>Ada.S | XP_018090401.1<br>NP_001085740.1 | 17336388<br>950506 | | No Ada activity in S.c. | | | | Adenosine deaminase |
| Ada2.L<br>Ada2.S | NP_001090531.1<br>NP_001089165 | 6254244<br>6251688 | | No Ada activity in S.c. | | | | Adenosine deaminase |
| Ada.2.S | NP_001087740.1 | 5929096 | ADA.2 | No Ada activity in S.c. | | | | Adenosine deaminase |
| Adk.S | NP_001086357.1 | 997231 | Ado1 | NP_012639.1 | 40 | *YJR105W* | 91 | Adenosine kinase |
| Adsl.L | NP_001080593 | 380285 | Ade13 | NP_013463.1 | 64 | *YLR359W* | 92 | Adenylosuccinate lyase |
| Adssl1.L<br>Adssl1.S | NP_001090012.1<br>NP_001087505.1 | 5758296<br>6254009 | Ade12 | NP_014179.1 | 57<br>57 | *YNL220W* | 92<br>92 | Adenylosuccinate synthase |
| Adss2.L | NP_001080088.1 | 944129 | Ade12 | NP_014179.1 | 56 | *YNL220W* | 92 | Adenylosuccinate synthase |
| Ak1.L<br>Ak1.S | NP_001087683.1<br>NP_001085451.1 | 6253770<br>6251725 | Adk1 | NP_010512.1 | 28<br>28 | *YDR226W* | 84<br>92 | Adenylate kinase |
| Ak2.L<br>Ak2.S | XP_018102289.1<br>NP_001080232.1 | 17344313<br>6254479 | Adk1 | NP_010512.1 | 55<br>59 | *YDR226W* | 94<br>90 | Adenylate kinase |
| Ak3.L<br>Ak3.S | NP_001089446.1<br>NP_001084561.1 | 977031<br>17332462 | Adk2 | NP_011097.3 | 46<br>45 | *YER170W* | 86<br>86 | Adenylate kinase |
| Ak4.L | XP_018113718.1 | 17341782 | Adk2 | NP_011097.3 | 43 | *YER170W* | 83 | Adenylate kinase |

| | | | | | | | | |
|---|---|---|---|---|---|---|---|---|
| Ak4.S | XP_018116248.1 | 959957 | | | 46 | | 83 | |
| Ak5.L<br>Ak5.S | XP_018113675.1<br>XP_018116212.1 | 6487811<br>17345311 | Adk2 | NP_011097.3 | 27<br>25 | *YER170W* | 61<br>58 | Adenylate kinase |
| Ak6.L<br>Ak6.S | NP_001089528.1<br>NP_001087040.1 | 972297<br>17335043 | | none | | | | Adenylate kinase |
| Ak7.S | NP_001081046.1 | 953552 | Adk1 | NP_010512.1 | 29 | *YDR226W* | 16 | Adenylate kinase |
| Ak8.L | NP_001088862.1 | 5831351 | | none | | | | Adenylate kinase |
| Ak9.L<br>Ak9.S | XP_018118807<br>XP_018118808 | 17335922<br>17335923 | Adk1 | NP_010512.1 | 24<br>24 | *YDR226W* | 14<br>14 | Adenylate kinase |
| Ampd1.L<br>Ampd1.S | XP_018101841.1\|<br>XP_018104675.1 | 17335636<br>17335637 | Amd1 | NP_013677.1 | 51<br>51 | *YML035C* | 68<br>68 | Adenosine monophosphate deaminase |
| Ampd2.S | XP_018104829 | 6489067 | Amd1 | NP_013677.1 | 53 | *YML035C* | 82 | Adenosine monophosphate deaminase |
| Ampd3.L<br>Ampd3.S | XP_018112762.1<br>XP_018115800.1\| | 6489056<br>17344991 | Amd1 | NP_013677.1 | 55<br>48 | *YML035C* | 64<br>80 | Adenosine monophosphate deaminase |
| Aprt.L | XP_018096991.1 | 6253243 | Apt1 | NP_013690.1 | 47 | *YML022W* | 93 | Adenine phosphoribosyltransferase |
| Atic.L<br>Atic.S | NP_001090100.1<br>XP_018094436.1 | 998812<br>17331460 | Ade17<br>Ade16<br>Ade17<br>Ade16 | NP_013839.1<br>NP_013128.1<br>NP_013839.1<br>NP_013128.1 | 62<br>62<br>61<br>62 | *YMR120C*<br>*YLR028C*<br>*YMR120C*<br>*YLR028C* | 99<br>99<br>99<br>99 | Amino-Imidazole CarboxAmide<br>Ribonucleotide transformylase<br>and IMP cyclohydrolase |
| Gda.L<br>Gda.S | NP_001083074.1<br>XP_018099342.1 | 987047<br>17337222 | Gud1 | NP_010043.1 | 39<br>39 | *YDL238C* | 96<br>91 | Guanine deamnase |
| Gart.L | NP_001093352.1 | 6251947 | Ade5,7<br>Ade8 | NP_011280.1<br>NP_010696.3 | 46<br>29 | *YGL234W*<br>*YDR408C* | 96<br>31 | Phosphoribosylglycinamide formyltransferase,<br>phosphoribosylglycinamide synthetase<br>and phosphoribosylaminoimidazole synthetase |
| Gmps.L<br>Gmps.S | XP_018119259.1<br>XP_018121231.1 | 950699<br>17340569 | Gua1 | NP_013944.1<br>NP_013944.1 | 37<br>36 | *YMR217W* | 93<br>94 | Guanine monophosphate synthase |

| | | | | | | | | |
|---|---|---|---|---|---|---|---|---|
| Guk1.L | NP_001087146.1 | 6254574 | Guk1 | NP_010742.1 | 55 | *YDR454C* | 92 | Guanylate kinase |
| Guk1.S | NP_001086807.1 | 17334557 | | | 55 | | 92 | |
| Hprt1.L | NP_001090235.1 | 6078713 | Hpt1 | NP_010687.3 | 34[a] | *YDR399W* | 20[a] | Hypoxanthine-Guanine phosphorybosyl transferase |
| Impdh1.L | | | Imd2 | NP_012088.3 | 60 | *YHR216W* | 97 | |
| | NP_001080792 | 957455 | Imd3 | NP_013536.3 | 63 | *YLR432W* | 97 | Inosine monophosphate dehydrogenase |
| | | | Imd4 | NP_013656.1 | 63 | *YML056C* | 97 | |
| IImpdh2.L | | | Imd2 | NP_012088.3 | 61 | *YHR216W* | 96 | |
| | NP_001083990.1 | 17345296 | Imd3 | NP_013536.3 | 64 | *YLR432W* | 96 | |
| | | | Imd4 | NP_013656.1 | 63 | *YML056C* | 98 | Inosine monophosphate dehydrogenase |
| IImpdh2.S | | | Imd2 | NP_012088.3 | 62 | *YHR216W* | 96 | |
| | NP_001082410.1 | 478732 | Imd3 | NP_013536.3 | 64 | *YLR432W* | 96 | |
| | | | Imd4 | NP_013656.1 | 63 | *YML056C* | 97 | |
| Paics.1.L | NP_001080163 | 17334793 | Ade1 | NP_009409.1 | 25 | *YAR015W* | 51 | Phosphoribosylaminoimidazole carboxylase |
| | | | Ade2 | NP_014771.3 | 31 | *YOR128C* | 29 | phosphoribosylaminoimidazolesuccinocarboxamide synthetase |
| Paics.1.S | NP_001086248.1 | 965998 | Ade1 | NP_009409.1 | 26 | *YAR015W* | 51 | |
| Paics.2.L | XP_018085590.1 | 5872998 | Ade2 | NP_014771.3 | 31 | *YOR128C* | 14 | Phosphoribosylaminoimidazole carboxylase phosphoribosylaminoimidazolesuccinocarboxamide synthetase |
| Pfas.S | XP_018094887.1 | 6485675 | Ade6 | NP_011575.1 | 37 | *YGR061C* | 93 | Phosphoribosylformylglycinamidine synthase |
| Pnp.L | NP_001079809.1X | 1011603 | Pnp1 | NP_013310.1 | 52 | *YLR209C* | 87 | Purine nucleoside phosphorylase |
| Pnp.S | P_018099448.1 | 17337633 | | | 45 | | 91 | |
| Ppat.L | NP_001083491.1 | 972209 | Ade4 | NP_014029.1 | 36 | *YMR300C* | 91 | Phosphoribosylpyrophosphate amidotransferase |
| Ppat.S | XP_041435416 | 17340761 | | | 36 | | 92 | |
| Xdh.L | XP_018117710.1 | 6485774 | | No Xdh activity in *S.cerevisiae*. | | | | Xanthine dehydrogenase/oxidase |
| Xdh.S | XP_018120131.1 | 17331587 | | | | | | |
